# Network analysis outlines strengths and weaknesses of emerging SARS-CoV-2 Spike variants

**DOI:** 10.1101/2021.09.03.458946

**Authors:** P.D. Manrique, S. Chakraborty, K. Nguyen, R. Mansbach, B. Korber, S. Gnanakaran

**Author notes:** Correspondence (SG).

## Abstract

The COVID-19 pandemic, caused by the SARS-CoV-2 virus, has triggered myriad efforts to dissect and understand the structure and dynamics of this complex pathogen. The Spike glycoprotein of SARS-CoV-2 has received special attention as it is the means by which the virus enters the human host cells. The N-terminal domain (NTD) is one of the targeted regions of the Spike protein for therapeutics and neutralizing antibodies against COVID-19. Though its function is not well-understood, the NTD is reported to acquire mutations and deletions that can accelerate the evolutionary adaptation of the virus driving antibody escape. Cellular processes are known to be regulated by complex interactions at the molecular level, which can be characterized by means of a graph representation facilitating the identification of key residues and critical communication pathways within the molecular complex. From extensive all-atom molecular dynamics simulations of the entire Spike for the wild-type and the dominant variant, we derive a weighted graph representation of the protein in two dominant conformations of the receptor-binding-domain; all-down and one-up. We implement graph theory techniques to characterize the relevance of specific residues at facilitating roles of communication and control, while uncovering key implications for fitness and adaptation. We find that many of the reported high-frequency mutations tend to occur away from the critical residues highlighted by our graph theory analysis, implying that these mutations tend to avoid targeting residues that are most critical for protein allosteric communication. We propose that these critical residues could be candidate targets for novel antibody therapeutics. In addition, our analysis provides quantitative insights of the critical role of the NTD and furin cleavage site and their wide-reaching influence over the protein at large. Many of our conclusions are supported by empirical evidence while others point the way towards crucial simulation-guided experiments.

## INTRODUCTION

The emergence and subsequent worldwide spread of the severe acute respiratory syndrome coronavirus 2 (SARS-CoV-2) causing COVID-19 [1, 2] is a global health emergency that, according to the World Health Organization, has taken more than 4 million lives as of July 16, 2021 [3]. COVID-19 is a highly contagious respiratory illness initiated by viral entry into host cells prior to infection and symptoms. The first event in viral entry is contingent upon the binding of the Spike glycoprotein, located at the surface of the SARS-CoV-2 pathogen, with the human host receptor angiotensin-converting enzyme 2 (ACE2) [4–6].

The Spike is a homotrimeric class I viral fusion protein able to adopt different conformations according to the state of the receptor-binding-domain (RBD) of each of its protomers. It has been determined by cryo-electron microscopy at the atomic level [7] that the Spike adopts two predominant conformations in which either all three protomers are in a closed state (all-down conformation), or one protomer is in an open state while the remaining two are in a closed state (one-up conformation). The one-up conformation promotes host receptor binding due to heightened exposure of the binding site region of the virus (the receptor binding domain or RBD). Each protomer of the Spike comprises around 1200 residues in the extracellular domain that can be grouped into two main functional subunits S1 and S2 delimited by the furin cleavage sites (FCS) loop at residues 675-690. The S1 subunit comprises the RBD that carries out the recognition and binding process to the ACE2 protein host in the human lung [8–10]. On the other hand, the S2 subunit manages viral-host-cell membrane fusion and subsequent viral entry. As we have done previously [11], we divide each subunit further for analysis, yielding a total of 12 domains for each protomer [Fig. 1].

**Figure 1:**
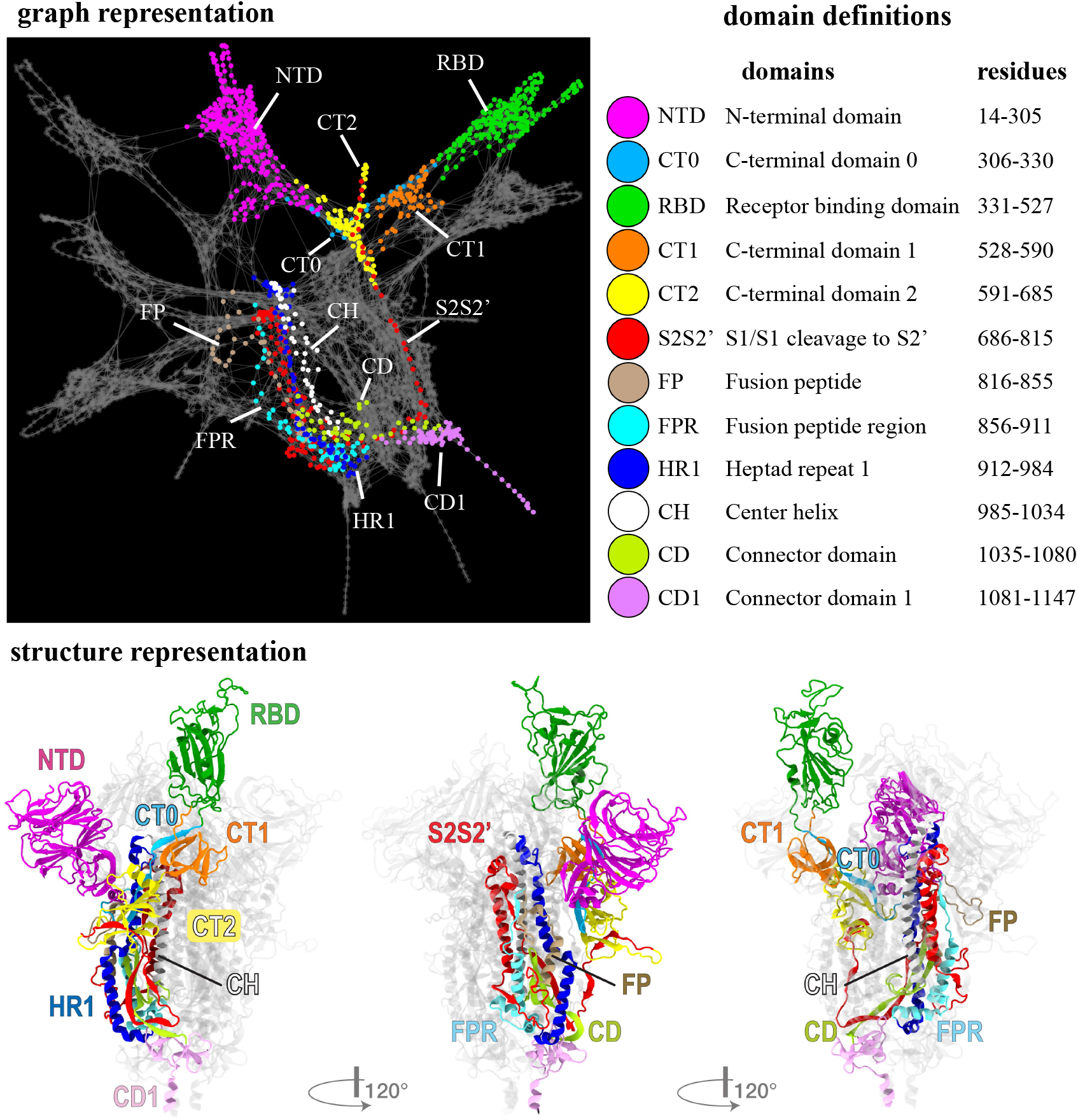
Graph and structure representations of the SARS-CoV-2 Spike protein. Top left and bottom panels illustrate the graph and graph representations, respectively, of the Spike protein divided into pre-assigned protomer domains (colored regions/nodes). In the top right panel, we provide the full domain names, the domain abbreviations, and the residues that comprise them.

From its discovery in November 2019 and throughout 2020, COVID-19, like other coronaviruses, has shown a slow rate of immunologically relevant variant accumulation, even after experiencing a worldwide spread, as compared to some other viruses such as influenza and HIV-1 [12, 13]. Indeed, the most notable evolutionary event has been on the Spike protein itself: a single amino acid substitution in residue 614 where an aspartic acid (D) was replaced by a glycine (G) [14]. This variant (G form) emerged at a significant level in early March of 2020 and quickly became the dominant form of the virus, holding an enhanced infection capability over the original (D form) [12, 14–17], by virtue of its greater RBD-open ACE2 binding capable conformation [17, 18]. Another potential factor impacting the infectivity of the virus is the higher rate and efficiency of furin cleavage found in the G form [19]. The Furin Cleavage Site (FCS) loop is an advantage acquired by the CoV-2 Spike over its CoV-1 predecessor, wherein furin-mediated S1-S2 proteolytic cleavage and conformational changes lead to more efficient downstream infectivity [20], and mutations near the furin cleavage sites may be contributing to enhanced infectivity of emerging variants [21, 22]. Most recently, several new and potentially significant variants have emerged, bringing with them additional challenges for vaccines and immunotherapies [23, 24]. Among them, recurrent deletions and accelerated substitution rates in the NTD drive antigenic and adaptive evolution and grant resistance to neutralizing antibodies [25, 26]. Most known NTD-specific neutralizing monoclonal antibodies (mAbs) target a specific region of the NTD, that coincides with the occurrence of these high-frequency mutations. This supersite that has emerged is composed of residues 14-20 (loop), 140-158 (*β*-hairpin), and 245-264 (loop) [27,28]. Though this mechanism of antibody escape has been minimized with NTD-binding cocktails of antibodies [29], it is still not quantitatively understood why the NTD is a crucial target for these binding antibodies and its relevance within the protein at large.

The functioning of the Spike protein is largely driven by allostery, with coupled dynamics between critical moving parts such as the NTD, RBD, FCS, and D614G-loop modulating its overall behavior. The manner by which these domains effectively communicate over long distances, however, remains unanswered. Here we implement a mathematical framework that sheds light on these puzzling questions and complements the current body of knowledge on the virus. We use extensive all-atom molecular dynamics simulations of the D and G forms in their dominant conformation states (all-RBD down and 1-RBD up) [11], to construct a weighted graph representation of the protein residues from the contact and correlation matrices of these simulations. With this framework, we quantitatively describe the dynamical relationships between different residues [30] and how they are impacted by the D614G substitution, as well as the inter-chain communication between either the three symmetrically down protomers, or the up-protomer with the left and right chains on either side (U, L, R-protomers respectively).

Using graph theory techniques, we identify the critical residues (nodes of the graph) and domains and assess their roles in communication and control of protein allostery. By engaging with these critical residues, we predict that it could drastically alter essential long-range molecular interactions and stability. Our analysis shows that these key residues tend not to overlap with those where most of the mutation/deletions occur. We hypothesize that the high mutation rate in regions such as the NTD occurs as a consequence of fitness pressures towards immune escape, while the lack of mutation in the key residues identified by the graph theory analysis demonstrates their likely importance for protein function. For calculations involving the RBD, we focus on the subset of residues that bind to the ACE2 receptor (residues 438-508) and hence are the most relevant for infectivity, the receptor-binding-motif (RBM). In specific, we find: (1) The communication structure in the G-form is more resilient to disruption than that of the D-form; (2) the G-form promotes efficient allosteric pathways to the RBM from distant regions such as the furin cleavage sites, at the cost of heightened vulnerability to RBD-binding antibodies compared to the D form; (3) enhanced symmetry in the use of hinge residues to communicate residue 614 with the receptor-binding-motif (RBM) in the one-up conformation of the G form over that of the D form, establishing more stable allosteric communication and robustness to eventual hinge mutations; (4) network measurements based on communication efficiency and node-to-node influence determine that key residues, most of them belonging to the NTD, are critically positioned and hierarchically connected to exert wide-reaching control of the protein at large; and (5) a specific examination of the NTD residues that are altered in the Delta variant, reveals that these residues are more efficient at impacting the full protein than the residues of the NTD supersite.

## RESULTS

We begin with the network characterization of the Spike protein. First, we address the differences between the closed and open conformations and how they are impacted by the D614G variant. We further illustrate the differences by looking into several aspects of the communication properties of each network, and how these differences impact infectivity and overall network stability. Due to abundance of theoretical and experimental data, this critical mutation serves as a solid test for the network approach. Then we move on to the importance of the functionally least understood NTD domain. Finally, we bring the network concepts to evaluate the emerging mutations and the critical sites for antibody binding. Additional network properties of the Spike protein, as well as, further details of the results presented in this section are provided in the Supplemenary Material (SI).

### Graph measure uncovers the communication core of the Spike protein and demonstrates that the D614G variant is more resilient to communication disruption

The ability of a particular node or group of nodes to form a bridge between distant regions of a network is quantified by the *betweenness centrality*. Mathematically, the betweenness centrality of a node is the number of optimal pathways (geodesics) that run through it as part of connecting any other pair of nodes in the network. In the context of our graph-based analysis of the Spike, residues taking part in a greater number of “shortest pathways”–i.e. the most highly-correlated series of contacts–connecting any other two residues of the protein are identified by higher betweenness centrality. Residues with high betweenness centrality act as critical ‘hubs’ that influence many optimal pathways of protein allosteric communication. We compute the betweenness centrality for each individual residue of our four networks and assess the implications at the level of the full protein, at the level of the domains, and at the level of the individual residues.

At the entire protein level, we find that the regions with the highest values of betweenness centrality form a closed ring about the protein, which forms the structural communication core of the whole system. Differences in the communication core between networks highlight the impact of the D614G shift in the all-down and one-up states [Fig.2(a)]. The ring straddles the equatorial plane of the Spike at the base of the NTD, extending via the CT0 to the base of the RBD. This high-centrality band of residues extends to the beta-strands at the nadir of the furin cleavage hairpin, and comprises residues of the NTD, CT0, CT1, and CT2 domains from all three protomers [Fig. S2]. This holds for both forms (D and G) and both conformations (all-down and one-up). However, we find that the communication ring in the G form is comprised of a greater fraction of residues of CT0 and a lower fraction of residues of CT2 and displays enhanced stability in the number of residues belonging to each of the regions when we compare the all-down and one-up conformations, in comparison to the D form. This core ring is crucial for communication between the three peripheral domains of the Spike - the NTD, RBD, and FCS (circled in black) and the central region of the protein. Interestingly, this high centrality core avoids the NTD supersite (highlighted in green in Fig. 2(a)), where many antibodies have shown particular binding preference and effectiveness [25, 27, 29]. The dynamics of all three domains involved in the core ring have been established to play significant roles in Spike protein function [31–33]. Our betweenness analysis therefore elucidates the allosteric linkage of these important moving parts of the Spike and demonstrates that a site of preferential antibody binding is not a site that impacts protein function.

**Figure 2:**
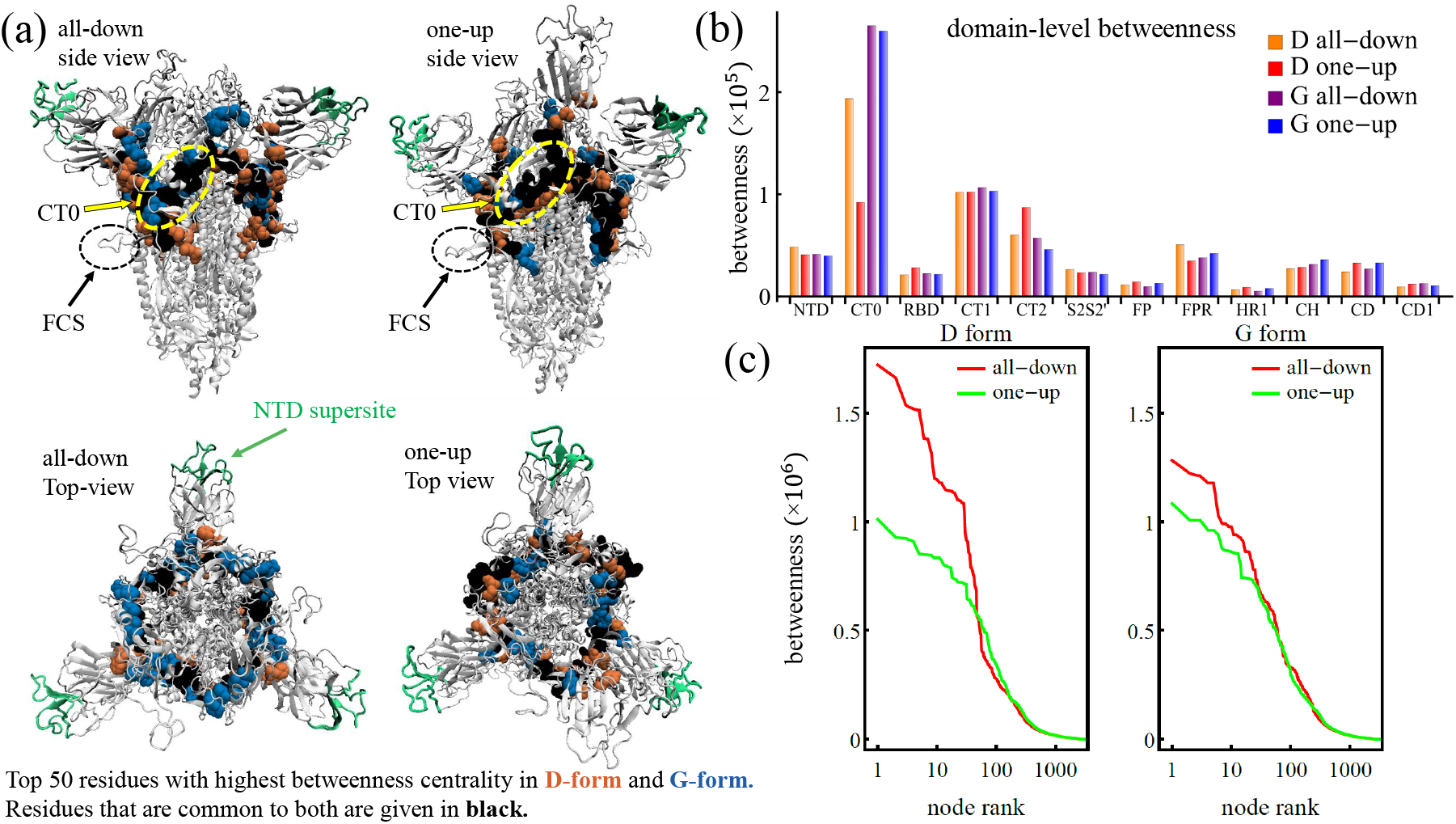
Communication core of the Spike protein. (a) SARS-CoV-2 Spike protein in the structure representation highlighting the NTD supersite (green), alongside the top-50 residues with highest betweenness centrality for the D form only (orange), the G form only(blue), and both forms (black), for the all-down (left panel) and one-up (right panel) conformations. (b) Betweenness centrality broken down by domain for the D and G form and the one-up and all-down conformations. (c) Ranked betweenness centrality for individual residues of the D form (left) and the G form (right) and the all-down (red) and one-up (green) conformations.

At the domain level, we find that CT0, despite being a small domain comprised of only 25 residues in each protomer [Fig. 1], is associated with the highest betweenness centrality overall. Indeed, it is the highest-ranking domain for both conformations of the G-form and the all-down conformation of the D-form (CT1 ranks slightly higher for the one-up D-form conformation) [Fig. 2(b)]. This domain, located at the heart of the Spike, appears to be crucial for communication between distant regions of the protein and hence could become a strategic target for intervention (i.e., drug target, antibody therapy). Our findings corroborate a previous experimental cryo-EM study of the Spike ectodomain [19] that speculated on the role of this NTD-RBD linker as a modulator of conformational changes connecting distal domains, and we go one step farther by using the centrality measures to quantify the relative significance of each residue in this role.

At the individual residue level, plotting the betweenness versus the node rank reveals that the all-down configurations contain nodes with higher values of betweenness, while the difference in betweenness for the two configurations is larger for the D-form than for the G-form [Fig. 2(c)]. In other words, the structural variation the Spike protein undergoes when transitioning from the all-down to the one-up configuration has a greater impact on its communication properties in the D-form than in the G-form. The G-form possesses more stable patterns of communication in which key functions of the protein should not be affected by the transition to the one-up state. This result agrees with the previously observed greater symmetrization of the inter-domain correlations and inter-protomer contacts in the G-form between the all-down and one-up states [11]. Moreover, from a purely networks perspective, the large values of betweenness in the all-down configuration of the D-form compared to those in the G-form uncover an increased vulnerability of the D-form, in which a small group of nodes are involved in a greater portion of the communication network. Perturbation of this group of nodes (e.g., through mutation, or binding partners) would more heavily influence the conformation of the Spike in the D form than a similar intervention performed on the G form, since a network with a more uniform distribution of betweenness centrality among its nodes tends to be more resilient to external interventions aimed at network disruption. This finding is in agreement with experimental and structural studies that found that the Spike protein of the D form is less stable with regards to S1 dissociation than that of the G form [15, 16].

### The G form promotes rapid allosteric pathways to the RBD at the cost of immune vulnerability

We explore the effects of the D614G variant at altering specific pathways relevant for viral infectivity through a path length analysis. Up to date, variants of concern are characterized by a combination of high transmissibility and/or immune response escape [23, 24, 26]. For example, the Delta variant, characterized by eleven modifications (two of them in the RBD) including two deletions, has shown resistance to monoclonal antibodies (mAbs) casirivimab and imdevimab, which are otherwise effective binders to the RBD able to inhibit human receptor binding [34]. Furthermore, RBD opening transition and changes in FCS are known to affect each other [18, 35]. Thus, variations in the communication properties to the RBD becomes a key question to address in order to establish the impact of mutations in the transmissibility and antibody resistance of the virus.

To this end, we analyze the optimal paths between the furin cleavage region and the RBM [Fig. 3(a)], and between residue 614 and the RBM [Fig. S4(a)]. We employ the Floyd-Warshall algorithm to determine the most efficient route connecting any two nodes in a given network from the contact matrix and the node pair correlations. The path length between these sites is defined as the sum of the edge weights of edges lying between the end point sites. As previously mentioned, high-magnitude correlations correspond to small edge weights and hence shorter paths, whereas low-magnitude correlations correspond to large edge weights and hence longer paths. Chains of highly correlated residues thus form shorter (i.e., more efficient) pathways than chains with the same number of links comprised by uncorrelated residues. A computation of the optimal pathways reveals that residue 614 lies near the shortest path from the FCS to the RBM region. This highlights its relationship to the up/down protomer states. Moreover, we find that the frequent mutation sites near FCS [36] do not generally lie on the optimal pathway to the RBM. We hypothesize that these sites do not overlap with the shortestpath sites due to the fact that changing the shortest-path sites would also change the communication pathway from the RBM to the cleavage site.

**Figure 3:**
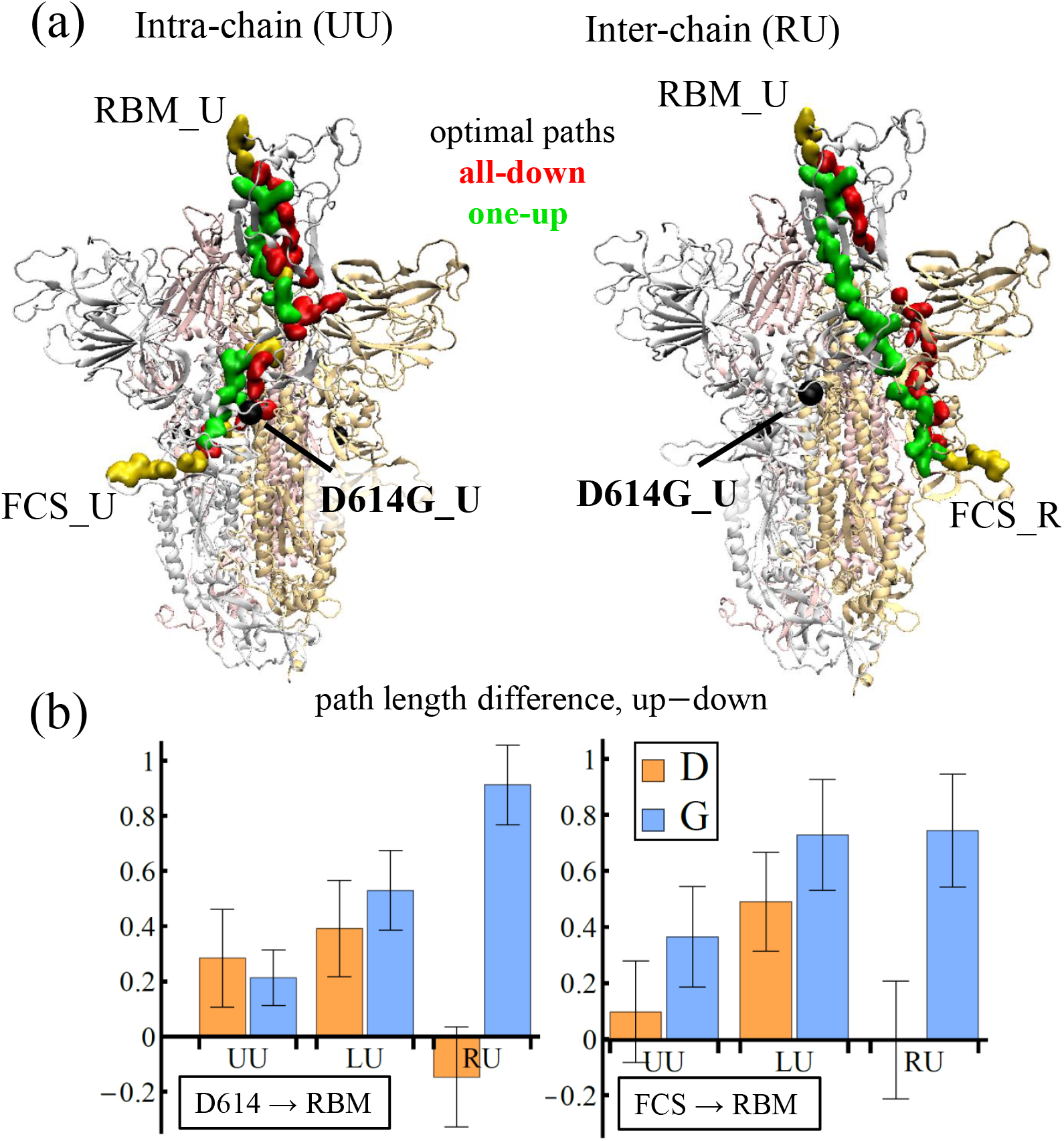
Optimal pathways to RBM from residue 614 and furin cleavage sites. (a) SARS-CoV-2 Spike protein in the structure representation highlighting the optimal intra-chain (left panel) and inter-chain (right panel) pathways from the furin cleavage (residue 685) to the RBM region (residue 501) for the all-down only (red), one-up only (green), and both conformations (yellow) in the G form. (b) Difference in the path lengths between the one-up and the all-down conformations of the D (orange) and G form (blue), for two sets of source/target regions: between residue 614 and the RBM (left panel) and between the furin cleavage site and the RBM (right panel). For all cases, the target region is specified at the protomer U, and the source is either protomer: from U to U (UU), from L to U (LU), and from R to U (RU).

In Figure 3(b), we depict the results of assessing optimal pathways between the different source/target regions. For consistency in the comparison between the two conformational states, we use the same protocol of focusing on the cases where the target chain is the U protomer, due to the up and down state variation of our simulations. Thus, each panel in Fig. 3(b) presents three pairs of results for each of the source/target options: from protomer U to protomer U (UU), from protomer L to protomer U (LU), and from protomer R to protomer U (RU). For the RBM and the furin regions, which are comprised of several residues, we compute the path lengths for all of the residues in the region and report the average distances alongside their fluctuations (error bar in the charts). Our results show that the G form experiences a larger disparity in the path lengths when comparing the all-down with the one-up conformation, where the all-down path length tends to be shorter than the one-up. Conversely, the D form shows smaller differences between the path lengths in these two conformations, and the all-down path lengths in the D form are generally longer than those of the G form [Fig. S4]. A rapid allosteric pathway to the RBM in the all-down conformation in principle reduces the effective RBD transition time to the up conformation that, in turn, enhances the overall binding effectiveness to the host receptor. The disparity tends to be enhanced in the inter-protomer pathways, of which, the counter-clockwise distances are more efficient [Fig. S4]. For most cases (five out of six), the path lengths to the RBM in the D form are longer than those of the G form. Hence, we could state that one of the mechanisms through which the D614G variant increases effective viral infectivity, is by enhancing the communication structure towards RBD while in the down conformation. Similarly, the length of the pathways in the up-conformation could be associated to the effective time elapsed to transition back to the down conformation, which becomes secondary within a viral infectivity context. However, a slow up-to-down transition of the RBD can be threatening to the virus as it presents an opportunity to bind to harmful agents such as antibodies. Therefore, our finding suggests that the G form enjoys viral infectivity advantages over the D form at the cost of a higher RBD vulnerability to antibodies. Additional details of these pathways and full list of residues involved are provided in Tables S4–S5 in the SI.

### Communication through the RBD hinge residues is more balanced in the G-form

The RBD is a globular domain that transitions between up (‘open’) and down (‘closed’) conformations (Fig. 1). Only in the open conformation is the ACE2 binding site appreciably exposed. This dynamic transition occurs via reorganization of two loops at the N-terminal and C-terminal bases of the RBD that act as “hinges” about which the RBD swivels. These two hinges H_1_ (residues 320-336) and H_2_ (residues 516-536) [Fig. 4(a)] are the only direct (covalently-linked) structural connections between the RBD and the rest of the Spike [5, 7] and therefore play a direct role in the regulation of the up and down movements of the RBD. Therefore, the perturbations at the distance stalk of the Spike, including the D614G shift, that have been shown to affect the gating mechanism of the RBD, must in some way communicate via the hinges [17, 18].

**Figure 4:**
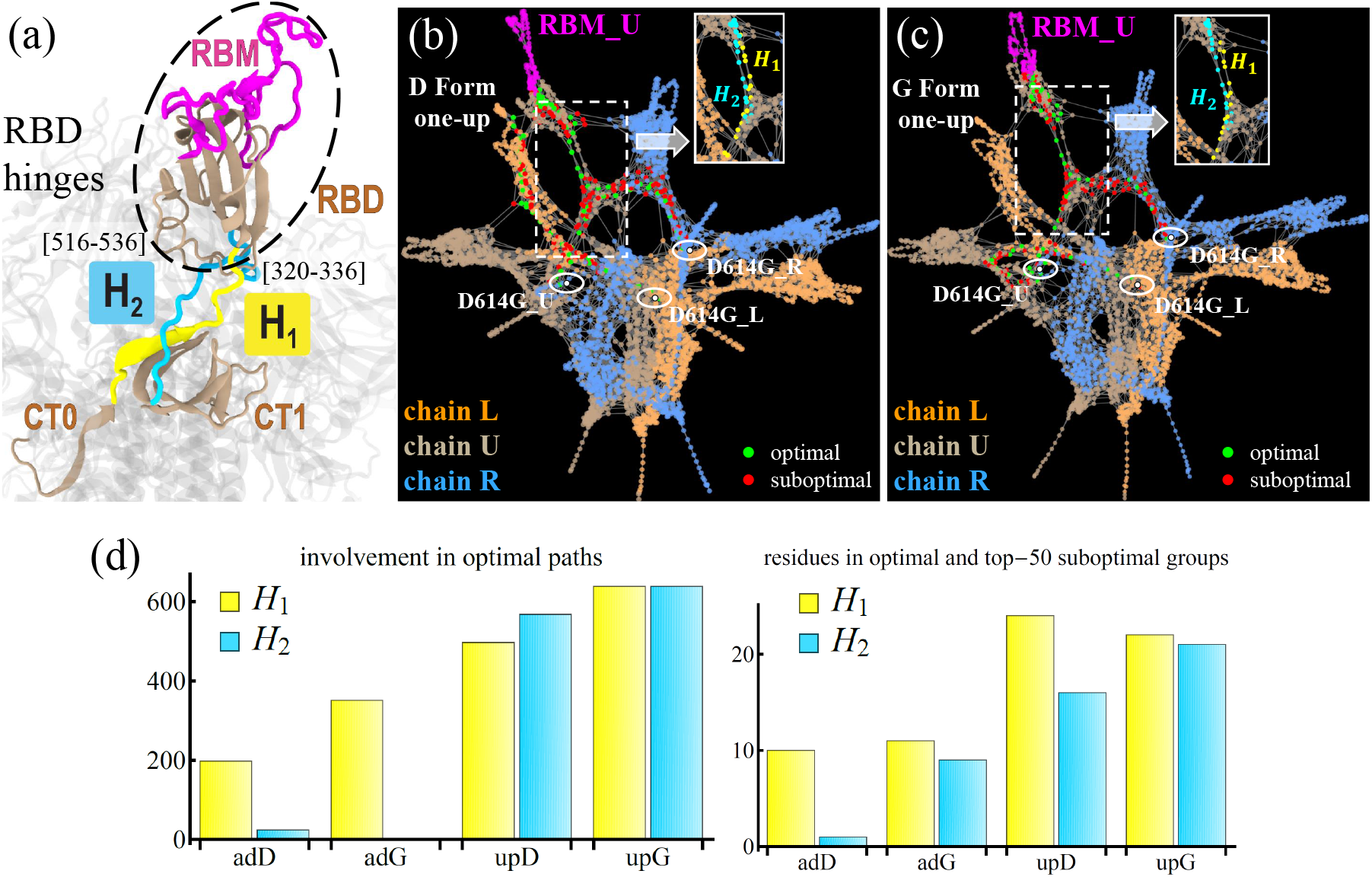
Hinge preferences in the D and G forms of the Spike protein. (a) Structural representation of a portion of the SARS-CoV-2 Spike protein highlighting the RBD hinges (H_1_ and H_2_) for the U protomer. (b-c) Spike protein in full in the graph representation for the one-up conformation of the D (b) and G (c) forms. Each network displays the protomers (chain) in a different color and highlights the RBM of the U protomer (magenta), as well as the residue 614 for each protomer (white nodes). Nodes comprising the optimal paths from 614 to RBM_U are colored in green, and the top-50 suboptimal nodes are shown in red (see text for the definition). Insets define the location of the hinge residues *H*_1_ (yellow) and *H*_2_ (cyan) in the network. (d) Number of times residues from H_1_ (yellow) and H_2_ (cyan) take part on the shortest paths from 614 to RBM U (left), and number of hinge residues that comprise the optimal paths and top-50 suboptimal nodes connecting residue 614 to RBM_U (right).

To understand how communication occurs between the stalk to the Spike, we explore the role of H_1_ and H_2_ in connecting residue 614 to the subset of residues of the RBD that binds to the ACE2, RBM (residues 438-508), by computing the shortest network pathways. We quantify the involvement of each of the hinge residues in these pathways in two complementary manners. In method (i) we count, for all RBM residues, the number of times a particular hinge residue is part of a shortest pathway between that residue and 614. In method (ii), we count the number of hinge residues that belong to either the group of residues comprising the shortest paths or an additional group comprising the top-50 suboptimal residues. Adding the top-50 suboptimal residues accounts for neighboring residues to the shortest pathway that could play a key role by assisting optimal residues in their communication tasks by providing alternatives routes to the RBM. These suboptimal residues are identified by comparing the direct (i.e., optimal) path from 614 and RBM with an indirect (i.e., suboptimal) pathway that goes through an additional third node, which does not belong to the optimal path. The difference in the length of these pathways provides a measure of how well this third node connects 614 and RBM through a suboptimal path; the smaller the path length difference, the higher the third node ranks within the suboptimal group. Thus, method (ii) identifies and quantifies the number of hinge residues that take part of optimal pathways and their surroundings (i.e., suboptimal pathways), and method (i) quantifies the frequency with which these residues are traversed in the optimal pathways.

Since the up conformation of the RBD is capable of binding ACE2, the event that leads to viral entry and later infection, we focus on the paths from residue 614 to RBM in the U-protomer [Fig. 4(b-c)]. In method (i), we examine a total of 852 paths, taking into account the relevant source/target combinations for the three protomers in all four protein configurations: D-form/G-form and all-down/one-up conformations. We find that in the all-down conformation there is a strong preference for using H_1_ over H_2_ irrespective of the D614G mutation. When the suboptimal residues are included, we observed that in the G form alone, the *H*_2_ residues began to participate in communication [Fig. 4(c)]. The D-form is found to avoid the use of the hinges in some of its inter-protomer paths. This is the case of R→U in the all-down conformation, and L→U in the one-up conformation [Fig. 4(b)]. The high inter-protomer correlations promote the formation of these low-cost alternative routes to RBM. Additional details of these calculations are provided in the SI.

The one-up conformation is characterized by an increase in the usage of *H*_2_ residues when compared to that in the all-down case. We find that the G form is able to use both hinges almost equally and at comparable frequencies. This is in contrast with the D form, where we find the use of a larger number of residues at a smaller frequency for H_1_ and a smaller number of H_2_ residues at larger frequencies [Fig. 4(d)]. Therefore, our findings show that, in the one-up state, the G form attains a greater balance in the usage of the hinge residues than the D form. With the G-form Spike known to have a higher opening probability [11,17,37], this greater balance establishes a more stable allosteric communication pathway, utilizing both the hinges and being more robust to eventual hinge mutations that may come up.

### NTD and CT are the communication hubs of the Spike protein

As its name suggests, closeness centrality measures the proximity of a particular node to the rest of the nodes in the network. Mathematically, it is defined as the inverse of the average path length between a particular node and all of the remaining nodes in the network. Higher values of this measure are associated with network-scale communication efficiency. This quantity answers the question of which nodes are most efficient at transmitting a signal (e.g., warning, cue, infection) to the rest of the network. In the context of the Spike protein, residues with a moderate to high closeness centrality are initiators of effective allosteric communication to able to efficiently reach every other residue of the protein.

Computing this centrality measure for each of our four graphs unveiled some commonalities, as well as, contrasting behavior at the protein, domain, and residue levels. At the protein level, in all four networks (D- and G-form, RBD-up and down), the S1 terminal dominates S2 in terms of high centrality [Fig. 5(a)]. Analyzing the residues that are within the top 33% in closeness, we find that a minimum of 98.76% and a maximum of 99.64% of them belong to S1, depending on the network. Though this is not necessarily surprising given the key role of the S1 subunit in regulating the receptor recognition/binding process, the finding provides a good check that validates the approach and provides additional support for our other findings. At the domain level, we find that the NTD, CT0, CT1, and CT2 regions score the highest in closeness centrality. This indicates that these domains are capable of drastically and efficiently impacting the entire protein. In the context of antibody neutralization, the virus is expected to mutate at regions where antibodies bind, for immune escape [26]. But if an antibody binds to the NTD, even if it is not exactly binding to the highest centrality residues, it will affect the NTD dynamics, which are highly correlated with the RBD dynamics, and therefore the antibodies are targeting areas of the protein that are highly relevant for protein function, possibly the up/down RBD transition.

**Figure 5:**
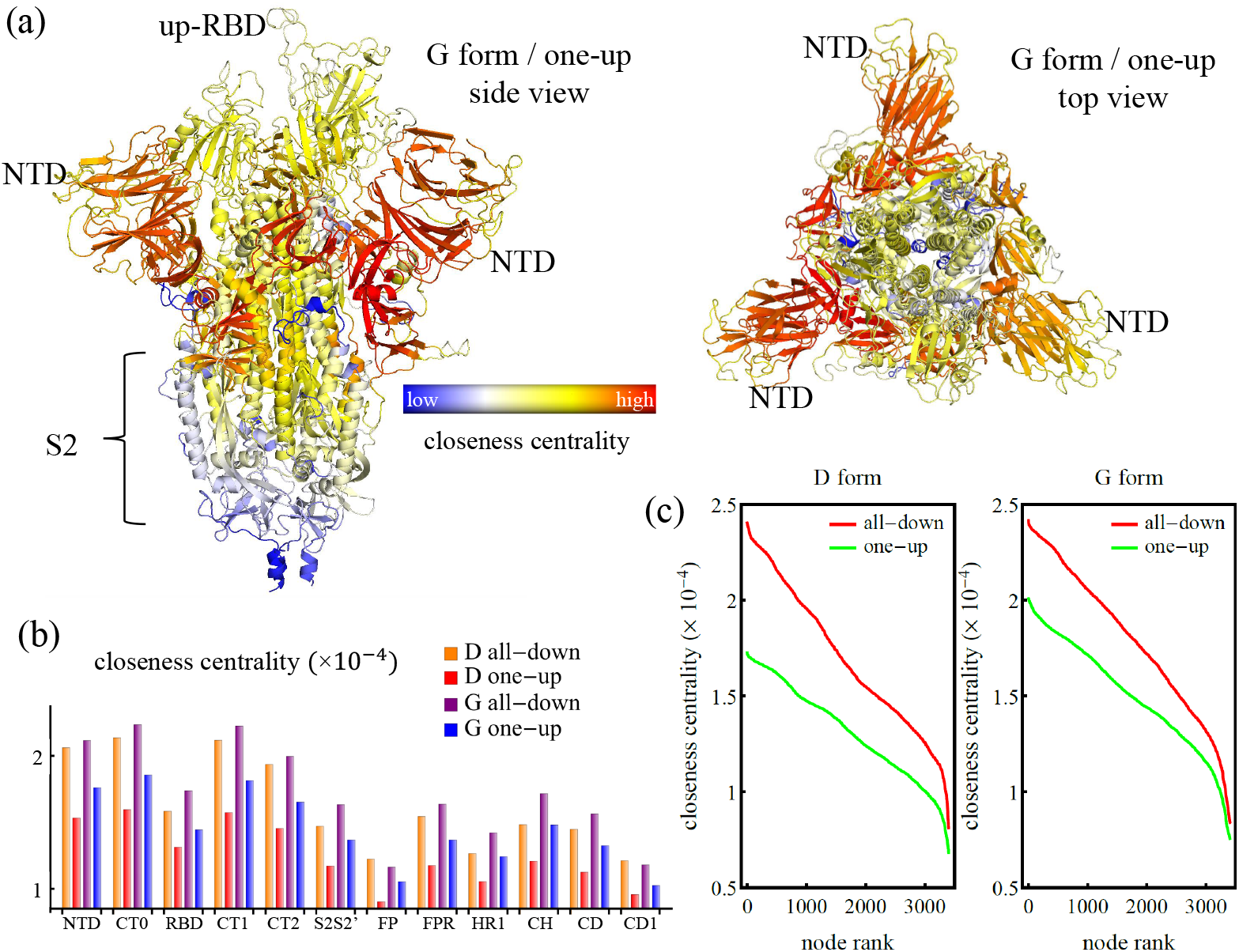
Closeness centrality highlights the NTD, CT0, CT1, and CT2 domains. (a) Structure representation of the SARS-CoV-2 Spike protein in the one-up conformation of the G forms colored by the closeness centrality. This measure is also presented broken down by protein region (b) and by individual residues ranked from highest to lowest (c), for the D and the G form, and for the all-down and the one-up conformations.

Comparing the closeness centrality of the different networks, a general trend is found where the closeness for the all-down conformation is higher than that of the one-up, and the closeness of the G-form is higher than that of the D form [Fig. 5(b)]. This statement also holds at the level of the individual residues [Fig. 5(c)]. A ranking of the nodes shows a smooth behavior in which, as opposed to the betweenness centrality of the top residues, the difference in closeness for consecutive ranked residues does not change drastically, and the overall trend of the closeness values of ranked residues agree with the domain-level trend. This tendency is explained by looking into the pair correlations of linked residues (i.e., edge weights) for the different protein systems. We find that the average pair correlation between contact residues for the all-down conformation in both, the D- and the G-form, is always higher than that of the one-up conformation, which in turn, makes the average path length shorter in the all-down conformation [Table S1]. Moreover, we find that the G-form has shorter path lengths, on average, than the D-form, which makes the Spike of the D614G variant more efficient at establishing effective allosteric communication pathways than that of the wild type.

### NTD is a central influence with a far-reaching role in protein functionality

While centralities of communication, such as betweenness and closeness, are focused on shortest paths analysis, centralities of influence, such as degree and eigenvector, take into account additional network features providing complementary information about the relevance of specific nodes at the local level as well as the network at large. Degree centrality quantifies the strength of the connections of a given node. For unweighted networks, this is simply the number of edges a particular node has. For weighted networks, each edge is scaled by its respective weight, and the sum of these weights is known as weighted degree or vertex strength [38]. Since it includes the nearest neighbors only, vertex strength is considered a measure of influence at the local level. An extension of this measure is eigenvector centrality, which computes the influence of a particular node based on the influence of its neighbors. Thus, the centrality score of a given node *j* takes into account the score of its neighbors, which in turn, take into account the scores of their neighbors, which are the second degree neighbors of node *j*, and so on. Therefore, this centrality intrinsically includes interactions with higher degree of neighbors, and hence quantifies the global influence of a given node. Mathematically, the eigenvector centrality of the node *j* is the *j*-th entry of the adjacency matrix eigenvector associated with the largest eigenvalue. In contrast to optimal paths analysis, in which edge weights indicate path lengths and therefore the smaller the weight the higher the importance of the edge, in influence centrality measures, the edge weights directly indicate importance. Therefore we employ an edge weighting for influence centrality that monotonically increases with the correlation between two residues (see Materials and Methods) and which is the inverse of the weight used for optimal communication paths calculations [39].

Computing the vertex strength and the eigenvector centrality of the graph representation of the Spike protein reveals that the NTD dominates in terms of both. At the domain level, a high vertex strength indicates that strong correlations between nearest neighbors in the NTD are common across the domain [Fig. S3]. This result is surprising: since the NTD is the largest domain of the Spike, containing 292 residues, one might expect a high number of pairs with lower correlations would drive down the average node strength. Other domains associated with strong intra-domain interactions are CT1, RBD, and CT0. In terms of eigenvector centrality, the NTD far outstrips any of the other domains in the protein, which highlights its hierarchical importance for the network at large [Fig. 6(a-c)]. This finding demonstrates in a quantitative manner that the critical residues in this domain, highlighted by the eigenvector centrality, exert wide-reaching influence to the protein at large, and that alteration in these nodes will affect the entire network. Interestingly, the residues with the highest eigenvector centrality in the G form are in close contact with the up-RBD but are not located within the NTD supersite [Fig. 6(a-b)]. It is likely that the experimentally-observed comparatively rapid mutation in the supersite occurs as a consequence of fitness pressures towards immune escape, while the lack of mutation in the nearby high eigenvector centrality residues further underscores their importance for protein function.

**Figure 6:**
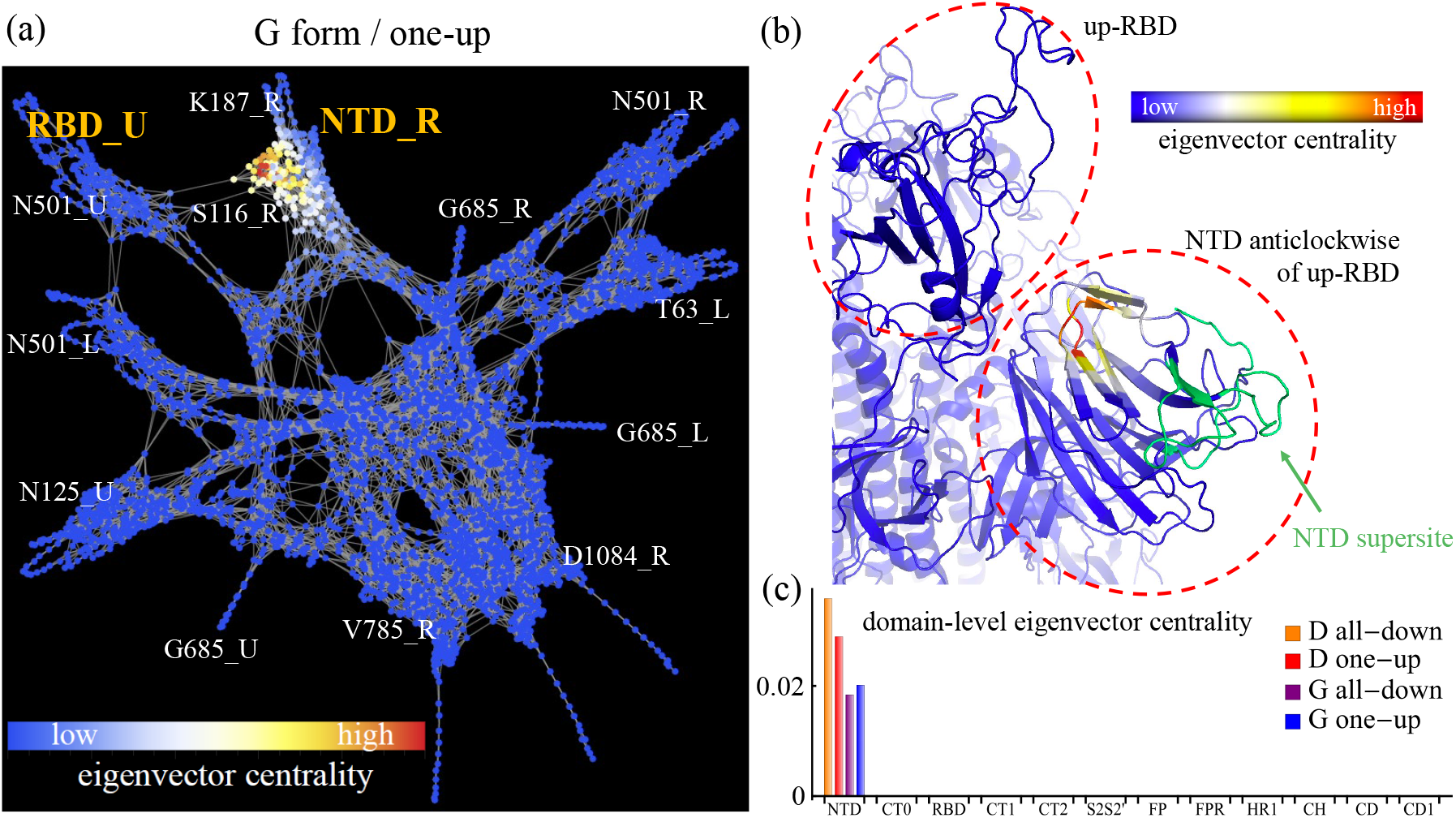
Eigenvector centrality highlights dominant role of NTD. (a) Graph representation of the SARS-CoV-2 Spike protein in the one-up conformation of the G form colored by individual node eigenvector centrality. (b) Structure close-up for the NTD and neighboring RBD region with eigenvector centrality highlighted by residue. (c) Domain-level eigenvector centrality for the D and the G forms, and for the all-down and the one-up conformations.

### NTD sites altered by the Delta variant impact the protein more efficiently than do the NTD supersite

Nine months after the emergence of the D614G variant in March of 2020, additional variants gradually emerged in several countries at different points in time [40–42]. Among the variants observed through July of 2021, the Delta variant (B.1.617.2), which originated in India [43], has received special attention due its enhanced infectivity, its ability to bypass antibodies [44], and to quickly become the dominant form of the virus, in a manner reminiscent of the previous year’s emergence of the D614G variant. Estimated proportions indicate that the Delta variant accounts for 57.6% of new infections in the US as of July 2021 and 98% of sequenced viruses circulating in the UK up to June 30, 2021 [45, 46]. The Delta variant possesses, in addition the D614G amino acid shift, three modifications and two deletions in the NTD supersite (T19R, G142D, R158G, and E156, E157, respectively), two modifications in the RBM (L452R, T478K), one modification near the FCS (P681R), and two additional modifications: one in the NTD (T95I) and another in the Heptad Repeat 1 (HR1) (D950N). The location of these sites in the Spike protein are shown in Fig. 7(a-b).

**Figure 7:**
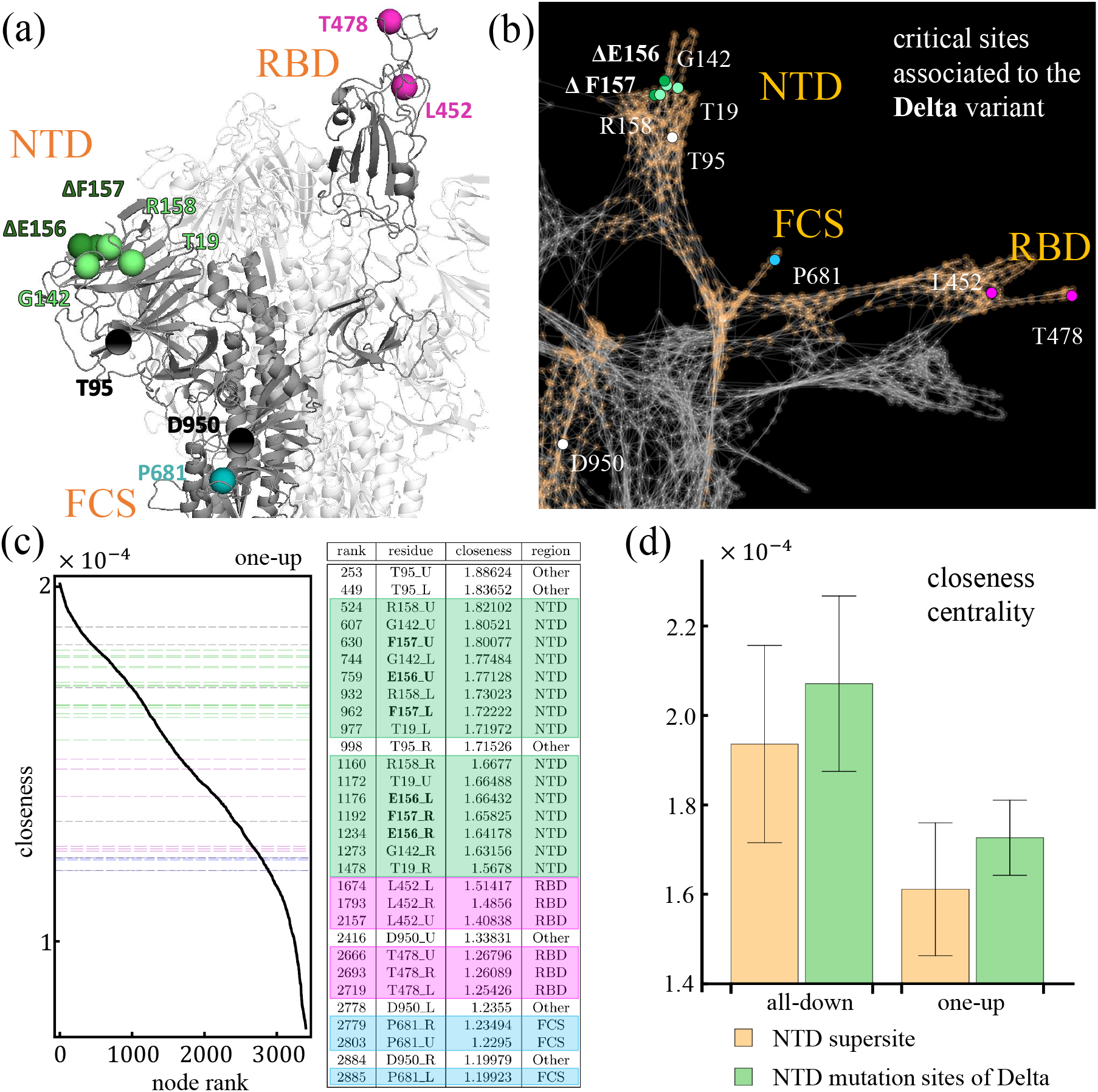
Relevance of sites modified/deleted by the Delta variant. (a) Structural representation of the SARS-CoV-2 Spike protein in the one-up conformation of the G-form highlighting the residues altered in the Delta variant. Deletions are marked in bold font and the Δ symbol. (b) Analogous to (a) but in the graph representation of the Spike protein. (c) Left panel shows the closeness centrality of individual residues ranked from highest to lowest in the one-up conformation of the Spike protein. The horizontal grid lines mark the values of closeness associated to the residues modified/deleted by the Delta variant. The colors match the table in the right panel, and shows detailed information concerning the individual ranking position, the identity of the residue (bold font indicates deletions), and the value of closeness in units of 10^−4^. (d) Average closeness centrality and fluctuations (error bars) of the NTD mutation sites of the Delta variant compared to that of the NTD supersite for the all-down and the one-up conformation.

We characterize the sites modified by the Delta variant with our network analysis, particularly the centrality measures of betweenness, closeness, and eigenvector. In terms of betweenness and eigenvector, we find that these residues rank very low for both conformations of the RBD, pointing to a scarce ability to bridge together or exert control over distant residues. Among the sites studied, the most relevant residue for communication (i.e., betweenness) is L452, through which runs a small fraction (5%) of the optimal communication pathways, in comparison to the highest ranked residue of the protein (V320 for all-down and V597 for one-up). As a background, residue 452 was extensively characterized as part of the California variant, and it impacted infectivity and resistance [47]. Deletions in this variant account for nearly 0.5% of the number of pathways carried by these highest ranked residues. Similar values are found when we examine the betweenness centrality of the modification at the FCS, and when we look at the eigenvector centrality of all the mutations [Fig. S5]. Therefore, modification/deletion of these residues are not expected to severely affect the protein in connectivity or large-scale control.

By contrast, examining the closeness centrality of these critical residues modified by the Delta variant, we find that those of the NTD rank high in their centrality measures (see Fig. 7(c) and Fig S6(a)) compared to the rest of the residues in the Spike. This means that these residues are good at initiating efficient pathways capable of reaching every region of the protein. They possess little capacity to serve as communication bridges (i.e., low betweenness), but they possess outstanding capacity to serve as pathway initiators (i.e., high closeness). These NTD residues effectively reach every other residue in the protein, with efficiencies ranging between 80-95% with respect to the highest ranked residue in closeness centrality (V597 for all-down, and K310 for one-up). Figure 7(d) compares the average closeness centrality for these sites with that of the residues comprising the NTD supersite (residues 14-20, 140-158, 242-264). We find that the average closeness of the NTD residues modified/deleted in the Delta variant is higher than the average closeness of the NTD supersite overall.

Our analysis provides a quantitative argument for why these critical sites are being modified/deleted in the Delta variant: changing residues with low betweenness does little to hinder the overall allosteric dynamics of the protein, but changing residues with high closeness could impact antibody binding. Modifications in the sites G142 and R158 in specific have already been shown to be directly associated to mAbs neutralization escape [27]. Thus, our results suggest that the closeness centrality may be used as a quantitative measure to determine critical residues in the Spike protein with the potential of serving as effective epitopes to neutralize the protein.

## DISCUSSION

The use of graph-derived approaches from MD simulations has been critical in the context of deciphering complex molecular interactions and their roles in regulating cellular processes [30, 48–50]. Many of these efforts have focused on the identification of the critical residues that facilitate effective allosteric interactions, how they are impacted by different conformational states, and how their modification could affect communication pathways [30]. It is thus an approach capable of guiding efforts aimed at the disruption of coordinated motion between distant and important functional regions in specific proteins.

In this article, we have implemented an approach of this nature to identify and analyze the critical residues pertaining to communication efficiency and wide-reaching control in the SARS-CoV-2 Spike protein. We studied a total of 20 *μ*s of all-atom MD simulations of the full protein trimer in the two widely-observed dominant conformations of the wild type as well as the dominant D614G variant. Our results are consistent with previous empirical and computational findings pertaining to the enhanced infectivity of the virus [12, 14, 16], the greater vulnerability to RBD-binding antibodies [17, 51], and the increased stability of the Spike protein, as a result of the D614G mutation [15, 16].

Long distance interactions and linkages in proteins are challenging to capture experimentally. For example, the mutational perturbation from D to G at position 614 is known to increase the occurrence of open Spike conformation [52] and enhance protease cleavage at FCS [53], but the exact mechanism of such information transfer has not been determined. Our computational graph approach has elucidated optimal pathways by which such distant sites of the Spike can communicate with each other and ultimately lead to concerted dynamics and allosteric modulations. Moreover, previous studies have highlighted the importance of hydrogen-bonds and structural reorganization of the RBD linkers acting as hinges [53, 54], and in this study we have established how critical communication through these hinges is more balanced in the D614G variant.

While some previous studies have implemented similar network-based approaches to on relatively shorter simulations with the aim of identifying overall subdomains and hubs in the Spike [55–57], our analysis provides detailed and new insights into the communication structure and the control cores of the protein at large. We identify a critical communication ring connecting peripheral regions of the protein, such as the NTD supersite, the up-RBD, and the furin cleavage sites, to each other as well as to the core of the protein, where crucial processes take place upon host receptor binding (e.g. membrane fusion). The structural ring is comprised of residues of the NTD, CT0, CT1, and CT2 domains. These regions also showed a remarkable ability to impact the entire network, where pathways initiated in these regions are able to reach every corner of the protein more efficiently. This provides a quantitative interpretation as to why there is a preference of some mAbs to bind to regions such as the NTD, and highlights the CT0, CT1, and CT2 domains as potential targets for functionality disruption and protein inhibition. Equally important, regions of the NTD were also highlighted by the eigenvector centrality, which is a control measure. The functional region found by our analysis does not overlap with the NTD supersite, or more generally, with high-frequency mutable residues [Table S6], suggesting that substitution/deletion events tend to avoid sites that are relevant for protein functionality. The dominant region is adjacent and in contact with the up-RBD pointing to a potential role of the NTD at facilitating RBD binding or initial virus attachment to the host-cell surface via recognition of specific sugar molecules, or possibly aiding the prefusion-to-postfusion transition, as found in other coronaviruses [58–60].

A specific examination of the NTD residues modified/deleted by the Delta variant according with these graph theory techniques, reveals that these critical residues are associated with an on-average greater closeness centrality than those of the NTD supersite, providing a quantitative explanation as to why these particular residues are been targeted for mutation, which adds to the factor associated to antibody resistance. Our analysis did not give a clear signal in the centrality measures associated to the modifications in the RBD of the Delta variant [Fig. S6]. However, examining the closeness centrality of the NTD residues modified/deleted in the Alpha, Beta, and Gamma variants, and comparing them with the closeness of the NTD supersite, a similar output as Fig. 7(d) is found for these variants [Fig. S7-S8]. This indicates that critical sites of the NTD, whether for mutation and/or antibody binding, can be characterized by their centrality values. With this in mind, we further examined the average centrality scores of all the residues in the G-form and find that a combination of lowbetweenness/low-eigenvector with mid-closeness and high-exposure, captures 80.4% of the residues in the NTD supersite, as well as, 63.1% of the NTD mutation sites found in the Alpha, Beta, Gamma, and Delta variants [Fig. S9]. Finally, we inspected residues 50 and 417 from the NTD and RBD, respectively, that are involved in a potential recombination event from pangolin RBD to the RBD found in SARS-CoV-2 leading to a significantly higher open conformation in CoV-2 as compared to earlier pangolin Spike [61]. While these two mutations can thus enhance the close-open transition, by virtue of local structural changes around residue 50 and reduced inter-RBD interactions at 417 [61], our network analysis indicates that the combination of relatively high closeness score and low betweenness score is maintained in this evolution as well. Specifically, residue 50 ranks in position 37 and 41 for the all-down and one-up conformation, respectively, in the D614G variant for closeness centrality. Their actual centrality values equal 98% of the highest-ranking node in closeness of each conformation. In the wild-type, the values are 96.5% and 97.8% of the highest ranked residue in the all-down and one-up conformations, respectively. The ranking of residue 417 in closeness is lower than residue 50 but still reaches scores within the 70% mark for all four networks. These residues rank below the 1.3% and 0.002% marks in betweenness and eigenvector centrality, respectively.

In summary, a network approach derived from all-atom MD simulations of the wild type and dominant variant of the SARS-CoV-2 Spike protein, provides quantitative insights related to the role of specific residues and regions for critical functions such as allosteric signaling, local- and global-level control, and network-scale communication efficiency. Our calculations identify the base of the NTD and the CT0, CT1, and CT2 domains as critical targets for communication disruption, and determine that a region of the NTD, adjacent and linked to the up-RBD, is able to exert wide-reaching influence over the whole system. Finally, a specific application of the employed network techniques to the NTD residues altered by the Delta variant points to a higher effectiveness of these residues at affecting the entire protein than that of the NTD supersite. Many of our conclusions are in agreement with empirical evidence while others point the way towards significant computational-guided experiments.

## MATERIALS AND METHODS

### Molecular modeling and simulation

For this work, we constructed a network model from the residue-wise correlation matrix of a series of extensive all-atom simulations we previously reported [11]. In brief, we constructed the initial structures based on experimentally-resolved Cryo-EM structures of the one-up and all-down states (PDB ID 6VXX and 6VYB). Regions–largely disordered loops– that were unresolved in these structures were built with a data-driven homology-modeling approach. Missing residues in the RBD specifically were built from an ACE2-bound RBD substructure (PDB ID 6M0J). The G-form was created from the D-form through manual mutation of residue 614, but otherwise the initial structures were identical. Atomistic molecular dynamics simulations were performed using the Gromacs software suite with the CHARMM36m force field. We ran five 1.2-*μ*s simulations for each of the four systems. The final 1000ns of each trajectory was considered as the production set, after equilibration, for a total of 20 *μ*s of simulation in neutral solution, using the Berendsen barostat, the particle mesh Ewald method for electrostatics and temperature coupling at 310K with a Langevin thermostat.

### Graph construction

We use the contact matrix in combination with the cross-correlation matrix to define the edges and weights of our networks, respectively. An edge between two residues is defined when heavy atoms of each residue were at a Euclidean distance of 6Å or less for at least 75% of the analyzed simulation frames. The cross-correlation matrix is defined as:

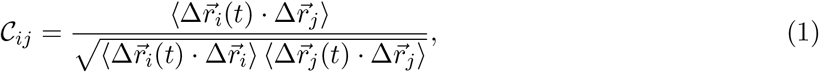

where 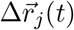 are the fluctuations of atom *j* with respect to its average coordinates. The weights for the communication graphs edges are defined as 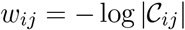 [30]. We use this definition of weights for calculations based on shortest paths analysis (e.g., betweenness and closeness centralities), while its inverse for control/influence analysis [39] (e.g. degree and eigenvector centralities).

### Definitions of network measures

In graph theory, centrality measures quantify the importance of a particular node within the network at large [62]. Since the importance can be interpreted in several different ways, there are different types of centrality measures according to specific criteria. In undirected networks, the most employed measures are degree, eigenvector, closeness, and betweenness centrality.

*Degree centrality* quantifies the number of edges a particular node has. It is interpreted as a measure of popularity and influence a node has at the local level. In weighted networks, each edge is leaden by a specific weight and the degree of a node is the sum of the weights of the associated edges and it is hence also known as *vertex strength*. Mathematically, the degree centrality of node *j* (*k_j_*) can be computed as:

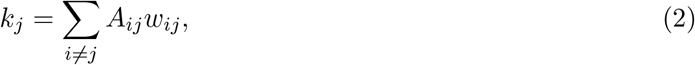

where *A_ij_* is the adjacency matrix defining the connections of the graph (*A_ij_* = 1, if *i* is in contact distance with *j* and *A_ij_* = 0, otherwise), and *w_ij_* is the weight matrix defining the strength in the connection between any two nodes.

*Eigenvector centrality* measures the influence a node has at a wider scale than degree. It quantifies the influence of a node given that of its neighbors. For example, a given node *j* has many edges and hence it would have a high degree centrality. But if its neighboring nodes have few or no connections, the influence of them is rather small and it will lead to a small eigenvector centrality for the node On the other hand, if the node *j* is connected with a few other nodes, it would hold a small value of degree centrality, but if these neighboring nodes are robustly connected, node *j* would hold an indirect influence over the additional nodes, which makes the eigenvector centrality of the node *j* higher. Thus, eigenvector centrality is a measure of influence on a larger scale. Mathematically, the eigenvector centrality of the node *j* is given by the *j*-th entry of the eigenvector 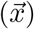 of the adjacency matrix (*A*) associated to its largest eigenvalue (*λ*). In closed form, the full eigenvector satisfies the following equation:

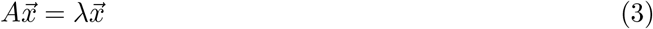

*Closeness centrality* measures how far a particular node is from all the other nodes in the network according to their associated pathlength, which quantifies the distance between any two nodes in a network. For weighted networks, the pathlength between node *i* and node *j* (*d_ij_*), is the sum of the weights associated to the edges that comprise the network path between node *i* and node *j*. Therefore, nodes holding higher values of closeness centrality are located, on average, at a shorter distance from all the rest of the nodes and hence are able spread information more efficiently throughout the graph. For a network with *n* number of nodes, the closeness centrality of the *j*-th node is defined as the inverse of the average distance (*l_j_*) between a particular node and the rest of the nodes.

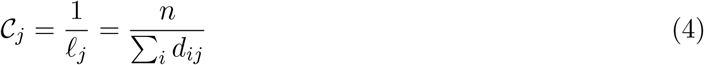

Finally, *betweenness centrality* quantifies the number of shortest pathways (also called geodesics) a particular node takes part on that connects any two other nodes. Hence, nodes with high betweenness are critical at establishing efficient bridges between distant regions of the network. The removal of these high-betweenness nodes can ultimately lead to the disruption of the network, and therefore their role is key for the stability and resiliency of it. Formally, betweenness centrality of node *j* (*b_j_*) in an arbitrary network can be expressed as:

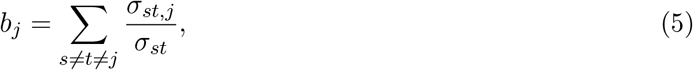

where *σ_st_* is the total number of geodesics connecting nodes *s* and *t*, while *σ_st,j_* is the number of geodesics connecting node *s* to node *t* that pass through node *j*.

## Acknowledgments

PDM and RM were supported by Los Alamos National Laboratory (LANL) Director’s Fellowship. SC was supported by the Center of Nonlinear Studies Postdoctoral program. KN and SG were partially supported by the DOE national laboratories focused on response to COVID-19, with funding providing by the Coronavirus CARES Act. BK and SG were partially supported by LANL LDRD project 20200706ER. This research used computational resources from LANL Institutional Computing Program. We thank Rory Henderson for insightful discussions.

## Contents

**A.** Spike network generalities

**B.** Communications core of the Spike protein

**C.** Shortest paths to receptor-binding-domain (RBD)

**D.** Involvement of the hinge residues of the up-RBD in shortest paths from 614

**E.** Impact of residue modification/deletions in the protein network

**F.** High frequency mutable residues

**G.** Harnessing centrality measures to characterize functionality regions in the N-terminal domain

### A. Spike network generalities

We construct the network representations of the Spike protein using the contact matrices and correlation of fluctuations associated to extensive all-atom molecular dynamics (MD) simulations of the D- and G-form, and in the all-down and one-up conformations of the receptor-binding-domain (RBD). An edge is established between residue *i* and residue *j* if the Euclidean distance of their heavy atoms are 6Å or less for at least 75% of the simulation frames. The undirected edges in our networks are weighted according to the correlations of pair of residues. Higher correlated residues (either positively or negatively) establish shorter path lengths than uncorrelated residues. The basic properties of the resultant networks are listed in Table S1. The first six columns list the number of intra- and inter-protomer edges, followed by the total number of edges (*m*), the average number of edges per node or mean degree 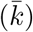, the average vertex strength 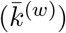, the average path length (*l*), and the unweighted and weighted clustering coefficients (C and C^(*w*)^, respectively). The latter two measures quantify how well are the neighbors of a particular node linked to each other. It is a measure of local cohesiveness of the network. An average over all nodes provides a global level of cohesiveness. Given that the weighted coefficient is greater than the unweighted one, indicates that interconnected triplets in the Spike protein are more likely formed by the edges with larger weights (higher correlations). In addition, the average path length points to stronger correlations in the all-down conformation than in the one-up, and also stronger in the G-Form than the D-Form. This result is echoed by the average vertex strength capturing higher values for the G-Form, as well. These contrasts are interesting and point directly to the dynamical effects, given that the topological values alone such as *m* and 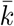, which do not account for the correlations, are very similar to each other for the four networks.

**Table S1:**
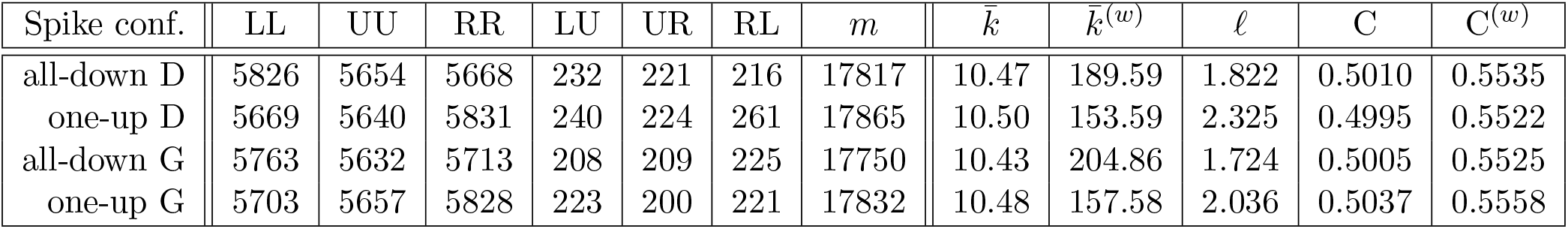
Basic network characteristics of the Spike protein. The properties outlined are: the number of intra-protomer edges (LL, UU, RR), inter-protomer edges (LU, UR, RL), total number of edges (*m*), mean unweighted degree 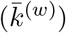, mean weighted degree 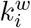, average pathlength (*l*), unweighted average clustering coefficient (C), weighted clustering coefficient (C^(*w*)^).

We further characterize the differences between the D- and G-Form of the Spike protein by looking at differences in the topology of the networks. A comparison between the adjacency matrices of the different networks provides the specific connections (i.e., edges) that are present in one network and absent in another one. Comparing the D- with the G-Form, we find that for the one-up conformation, a total of 676 edges are in the G-Form but not in the D-Form, while 709 are in the D-Form and absent in the G-Form. These values quantify the gain and loss edges, respectively, that the G-Form has over the D-Form. The net change is 1385 edges between G and D in the one-up conformation. In a like manner, the net change for the all-down conformation is 1445 edges, where 689 edges are gained, and 756 edges are lost. Figure S1 quantifies the excess edges (gain and loss), when we look at the specific domains/regions (see Fig. 1 of the main paper for the definition). As shown, for both conformations, the largest changes happen in the edges belonging to the NTD and the RBD. Fewer changes in the edges are found within the remaining regions, while no change occurs within the CT0. Looking at the inter-region changes, we find that the largest gain and loss in the all-down conformation occurs in the S2S2’ region, where the connection to the FPR is strengthened while the inter-region connection with the CH is weakened. By the same token, in the one-up conformation we find that the connections between CT0 and CT2, and between S2S2’ with FPR and with CH, are all weakened, while the connection between S2S2’ with HR1, is strengthened.

The results presented in the Table S1 were average values over the whole protein. Figure S3 details the region/domain level as well as the single-residue level behavior of the vertex strength and weighted clustering coefficient. As shown, the N-terminal domain (NTD) is dominant in the vertex strength measurement for all three protomers and all networks. This is in contrast with the clustering coefficient (Fig. S3(b)), where the values for most residues and all regions/domains are well balanced with each other. The weighted clustering coefficient for residue *i* is computed using Vespignani et. al. [1] expression for weighted networks:

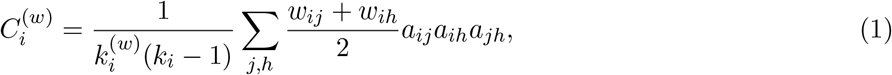

where 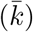 is the vertex strength of node *i*, *k_i_* is the unweighted degree, *w_ij_* is the edge weight connecting nodes *i* and *j*, and *a_ij_* is the adjacency matrix element associated to nodes *i* and *j*.

**Figure S1:**
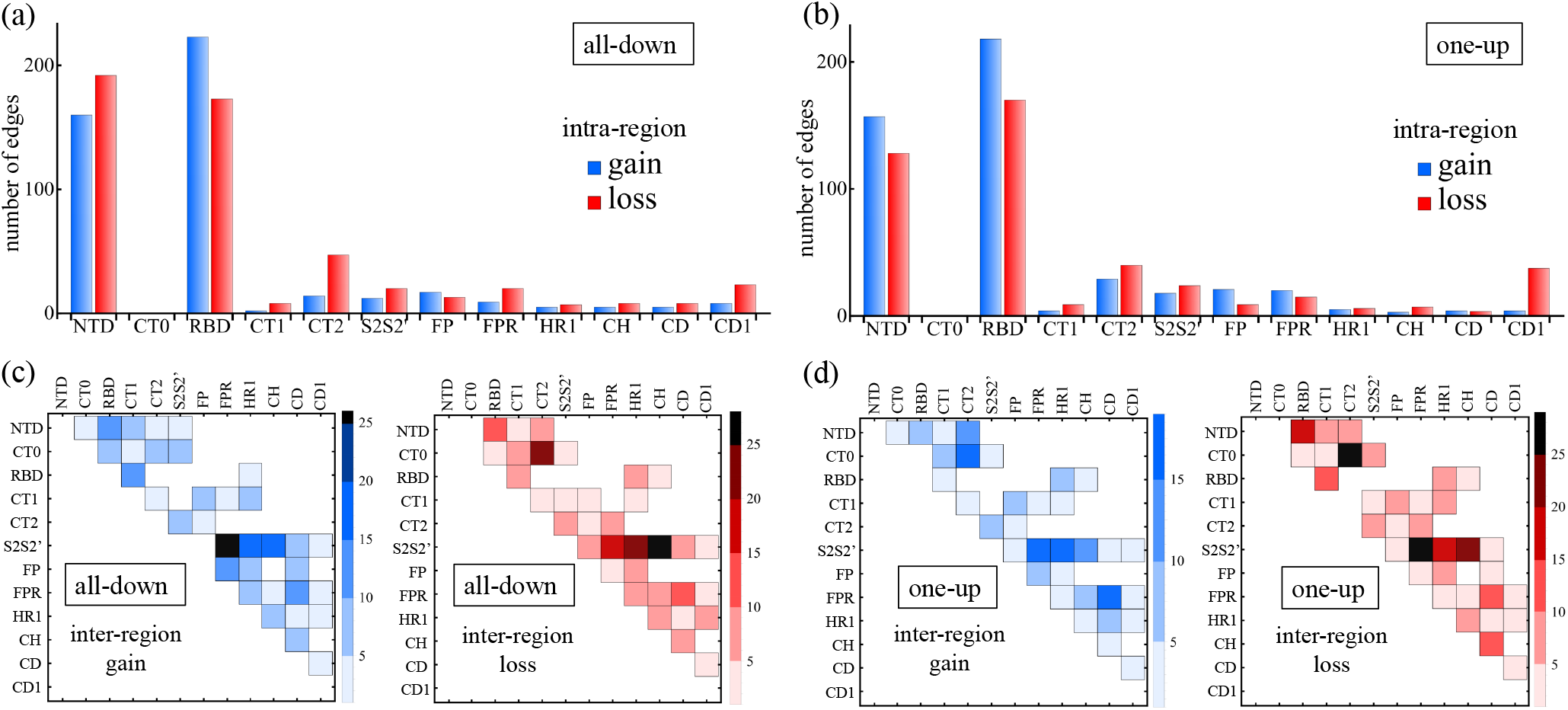
Comparison between the networks of the D- and G-Form using the excess of edges for the different regions/domains (see Fig. 1 for the definitions). Blue means a gain of edges of G over D while red means a loss of edges of G over D. (a-b) Intra-region edge differences. (c-d) Inter-region edge differences. The all-down conformation is shown in (a) and (c), while the one-up conformation in (b) and (d).

### B. Communication core of the Spike protein

Table S2 details the number of residues belonging to specific regions (see Fig. 1 of main paper, and Fig. S2) in the Spike for the top-50 ranking according to betweenness centrality. These residues comprise the communications ring depicted in the Figure 2 of the main draft. As shown, a greater participation of CT0 is found in the G-form, which is also characterized by an enhanced stability in the number of residues for each of the regions when we compare the all-down and one-up conformations. A detailed list of the top-50 residues ranked by betweenness centrality for the four networks is provided in the Table S3.

**Table S2:** Region/domain participation (number of residues involved) within the communication ring for the D- and G-form in the all-down and one-up conformations.

|  | NTD | CT1 | CT2 | CT0 |
| --- | --- | --- | --- | --- |
| D-form/all-down | 19 | 11 | 8 | 9 |
| D-form/one-up | 15 | 13 | 21 | 1 |
| G-form/all-down | 17 | 15 | 6 | 12 |
| G-form/one-up | 13 | 14 | 6 | 17 |

**Figure S2:**
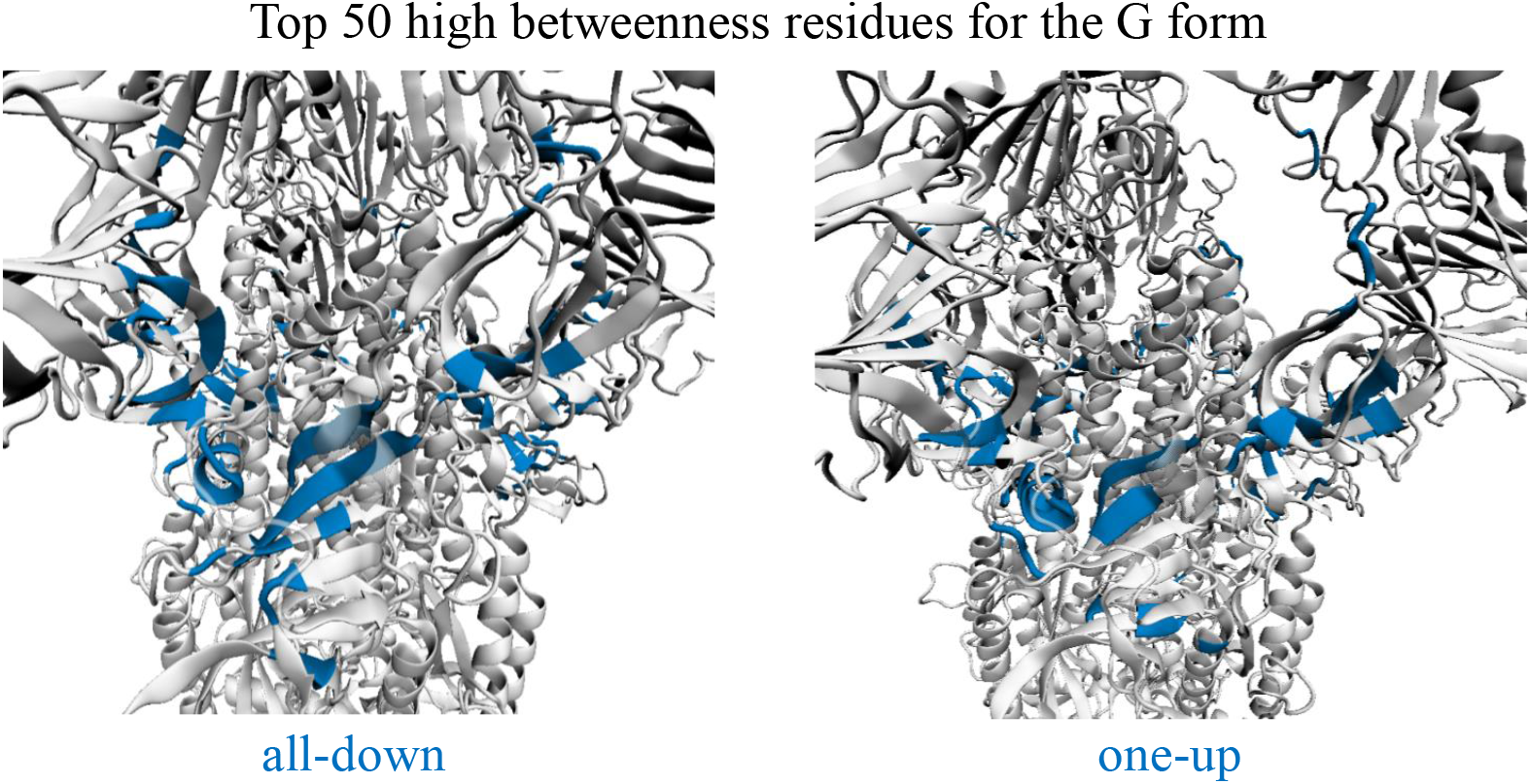
Top 50 residues with the highest values of betweenness centrality forming the communication core of the Spike protein. Illustration drawn for the G form in the one-up (left) and the all-down conformations (right)

**Figure S3:**
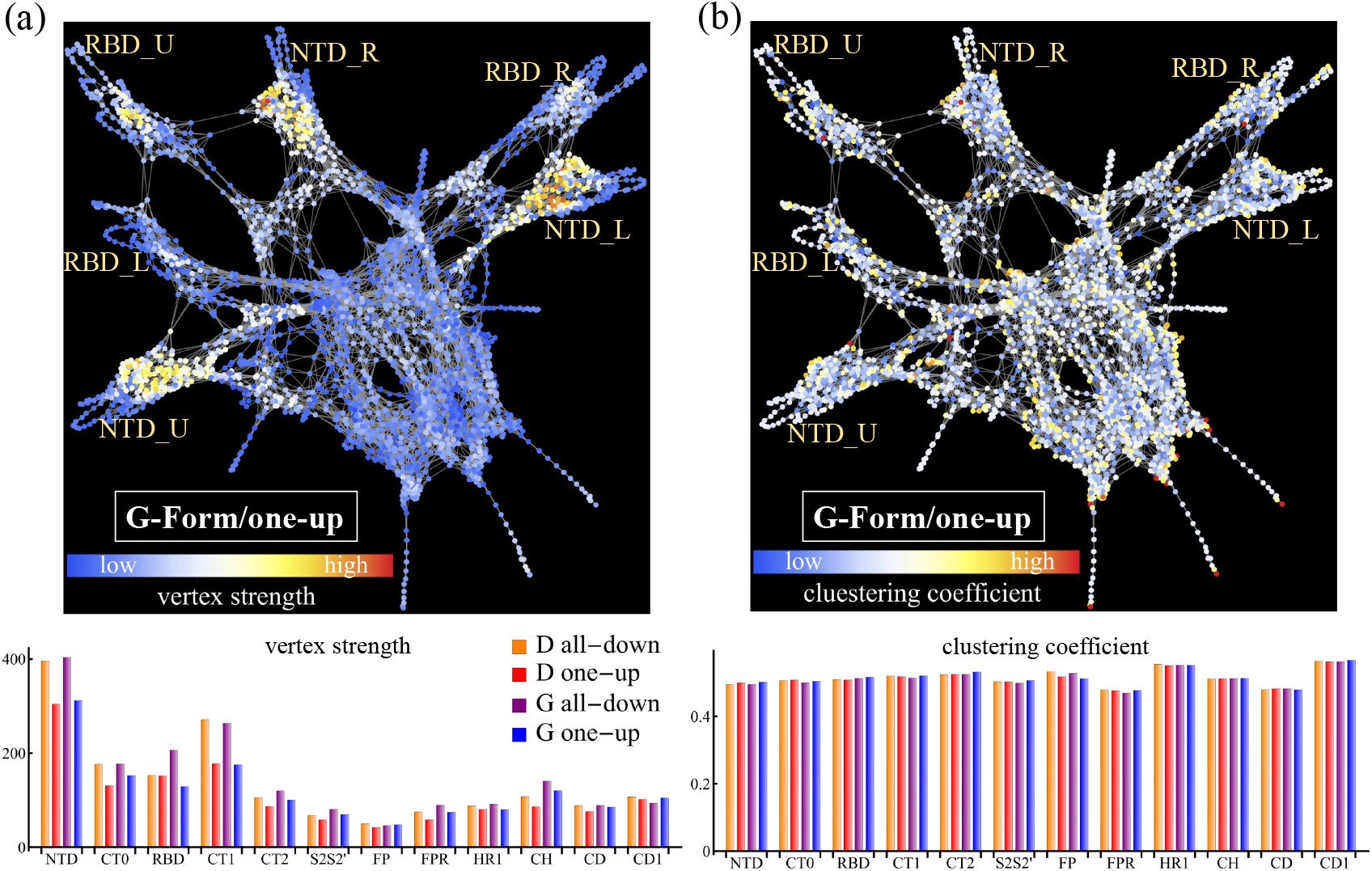
(a) Vertex strength in the one-up conformation of the G-Form in the network representation of the Spike protein (top panel), and vertex strength for the D- and G-Forms in the all-down and one-up conformations broken down by domains/regions (bottom). (b) Clustering coefficient in the one-up conformation of the G-Form in the network representation of the Spike protein (top panel), and clustering coefficients for the D- and G-Forms in the all-down and one-up conformations broken down by domains/regions (bottom).

**Table S3:** Top-50 residues ranked by betweenness centrality for our four networks (D- and G-form in the one-up and all-down conformations). Regions where the residues belong to as well as the values of betweenness centrality are also listed for reference.

| D-form/all-down |  |  | D-form/one-up |  |  | G-form/all-down |  |  | G-form/one-up |  |  |
| --- | --- | --- | --- | --- | --- | --- | --- | --- | --- | --- | --- |
| region | residue | betweenness | region | residue | betweenness | region | residue | betweenness | region | residue | betweenness |
| NTD | S297_U | $1.72 \times 10^6$ | CT2 | V597_L | $1.01 \times 10^6$ | CT0 | V320_R | $1.28 \times 10^6$ | CT2 | V597_L | $1.08 \times 10^6$ |
| NTD | L296_U | $1.66 \times 10^6$ | CT2 | V597_R | 928258 | CT0 | F318_R | $1.22 \times 10^6$ | NTD | S297_L | $1.01 \times 10^6$ |
| CT0 | V320_U | $1.53 \times 10^6$ | CT1 | G566_U | 922602 | CT0 | N317_R | $1.21 \times 10^6$ | NTD | L296_L | $1.01 \times 10^6$ |
| CT0 | R319_U | $1.52 \times 10^6$ | NTD | R44_R | 909094 | CT0 | T315_R | $1.18 \times 10^6$ | CT0 | V320_L | 961043 |
| CT0 | F318_U | $1.51 \times 10^6$ | CT2 | V595_L | 851495 | CT0 | S316_R | $1.18 \times 10^6$ | CT0 | R319_L | 960795 |
| NTD | Y279_U | $1.38 \times 10^6$ | CT2 | V595_U | 849469 | CT2 | V597_R | $1.03 \times 10^6$ | CT0 | F318_L | 941009 |
| NTD | R44_U | $1.38 \times 10^6$ | NTD | L296_L | 846836 | CT1 | L552_R | 990742 | CT1 | F565_L | 875319 |
| CT1 | G566_L | $1.32 \times 10^6$ | NTD | S297_L | 843956 | CT1 | V551_R | 990594 | CT0 | F318_U | 866657 |
| CT1 | G566_U | $1.2 \times 10^6$ | CT2 | G593_L | 833754 | CT1 | C538_R | 977307 | CT1 | V576_L | 865061 |
| CT1 | A575_L | $1.19 \times 10^6$ | CT2 | F592_L | 833622 | NTD | R44_U | 976806 | CT0 | V320_U | 861754 |
| CT1 | A575_U | $1.18 \times 10^6$ | CT2 | G594_L | 833423 | CT0 | R319_L | 940940 | CT0 | T315_U | 855051 |
| CT1 | I587_U | $1.18 \times 10^6$ | CT1 | C590_L | 820818 | CT0 | F318_L | 940562 | CT0 | S316_U | 853834 |
| CT1 | C538_U | $1.16 \times 10^6$ | CT2 | S596_U | 800920 | CT0 | V320_L | 932228 | CT0 | N317_U | 852772 |
| CT2 | V597_U | $1.15 \times 10^6$ | CT1 | A575_U | 797432 | CT0 | N317_L | 927872 | NTD | D290_L | 809650 |
| CT1 | V551_U | $1.14 \times 10^6$ | CT2 | V597_U | 794295 | CT1 | C538_L | 917349 | CT0 | V320_R | 741902 |
| CT2 | V595_U | $1.14 \times 10^6$ | CT2 | S596_L | 790348 | CT1 | G566_L | 904403 | CT0 | F318_R | 741118 |
| CT2 | S596_U | $1.14 \times 10^6$ | CT1 | I587_U | 789084 | CT2 | F592_U | 863330 | CT1 | V551_L | 741081 |
| CT1 | C538_L | $1.14 \times 10^6$ | CT1 | V576_L | 743093 | NTD | P295_R | 862415 | CT0 | R319_R | 738820 |
| CT1 | V551_L | $1.13 \times 10^6$ | CT1 | T588_U | 737910 | CT1 | C590_U | 862286 | CT1 | L552_L | 738663 |
| CT1 | L552_L | $1.13 \times 10^6$ | CT1 | C590_U | 735819 | NTD | D294_R | 860246 | CT1 | L585_L | 737992 |
| CT1 | D586_L | $1.11 \times 10^6$ | NTD | L276_U | 732330 | CT0 | V308_U | 831310 | CT2 | S596_L | 735785 |
| NTD | K278_U | $1.1 \times 10^6$ | NTD | Y279_R | 723910 | NTD | C291_R | 820395 | CT2 | V595_L | 734241 |
| CT0 | V320_L | $1.1 \times 10^6$ | NTD | K278_R | 721182 | CT2 | V595_U | 784027 | CT1 | C538_L | 728821 |
| NTD | V289_U | $1.1 \times 10^6$ | CT2 | S591_U | 720331 | CT2 | G593_U | 779075 | NTD | Y279_U | 718238 |
| NTD | A288_U | $1.1 \times 10^6$ | CT2 | F592_U | 720193 | CT2 | G594_U | 778918 | NTD | F43_U | 710466 |
| CT2 | V597_L | $1.09 \times 10^6$ | NTD | D290_L | 719393 | NTD | R44_L | 736157 | NTD | R44_U | 706687 |
| CT0 | R319_L | $1.09 \times 10^6$ | NTD | K300_U | 719390 | CT1 | G566_R | 731982 | NTD | P57_L | 696421 |
| CT0 | F318_L | $1.08 \times 10^6$ | CT2 | G594_U | 714378 | CT1 | A575_R | 701646 | NTD | L56_L | 696024 |
| NTD | P295_L | $1.04 \times 10^6$ | NTD | L276_R | 713872 | CT0 | F306_U | 696022 | CT2 | I664_U | 676359 |
| NTD | L276_R | 949065 | CT2 | G593_U | 713713 | NTD | Y279_U | 678263 | CT2 | I670_R | 654870 |
| NTD | L277_R | 916886 | NTD | L277_R | 712150 | NTD | K278_U | 675650 | CT2 | G669_R | 654435 |
| NTD | L293_L | 911165 | CT2 | C671_U | 646368 | CT1 | V551_U | 662846 | CT1 | S530_U | 636062 |
| NTD | D294_L | 908653 | CT2 | S591_R | 642849 | CT1 | L552_U | 661948 | CT1 | K528_U | 633144 |
| CT0 | V308_R | 858046 | CT1 | C590_R | 641995 | CT1 | D586_R | 656010 | CT1 | K529_U | 631971 |
| NTD | K300_R | 853131 | CT0 | R328_U | 640887 | NTD | T274_R | 645961 | NTD | L276_U | 618830 |
| NTD | F58_L | 847875 | CT2 | F592_R | 639899 | NTD | R273_R | 638207 | NTD | L277_U | 612963 |
| NTD | P57_L | 789889 | NTD | L277_U | 629095 | CT0 | T315_L | 633670 | CT0 | Y313_U | 611506 |
| NTD | L56_L | 789183 | CT1 | V576_U | 616462 | NTD | L270_R | 632052 | CT0 | Q314_U | 609766 |
| CT2 | V595_L | 780325 | CT1 | S530_U | 614864 | NTD | Q271_R | 631478 | NTD | K278_U | 599051 |
| CT2 | S596_L | 773370 | CT1 | K529_U | 612302 | NTD | C291_L | 630026 | CT1 | C538_U | 596039 |
| CT0 | K310_R | 752957 | CT2 | I670_U | 611603 | NTD | E298_L | 624214 | CT1 | V551_U | 585234 |
| CT0 | E309_R | 750877 | CT1 | K528_U | 611114 | CT1 | V551_L | 620519 | CT1 | L552_U | 584019 |
| CT2 | I664_R | 725737 | CT2 | V595_R | 594692 | CT1 | L552_L | 619554 | CT1 | V551_R | 551659 |
| NTD | R44_R | 702033 | CT2 | S596_R | 590924 | NTD | L276_R | 600233 | CT1 | C538_R | 551062 |
| NTD | S45_R | 695434 | CT1 | L585_L | 581346 | CT1 | A575_L | 599019 | CT0 | V308_U | 538623 |
| NTD | Y279_R | 682479 | NTD | R44_U | 575030 | NTD | Y279_R | 596696 | NTD | K300_U | 537585 |
| S2S2' | I692_U | 568037 | NTD | Y279_U | 571436 | NTD | R44_R | 593659 | CT0 | K310_R | 536400 |
| FPR | M869_R | 568011 | NTD | F43_U | 568408 | CT2 | I598_U | 590405 | CT0 | V308_R | 535633 |
| FPR | L864_L | 556447 | NTD | K278_U | 567140 | NTD | L277_R | 589463 | CT0 | E309_R | 535529 |
| CT2 | V608_U | 550331 | CT2 | G594_R | 562228 | CT1 | D586_L | 587652 | NTD | R44_R | 516465 |

### C. Shortest paths to receptor-binding-domain (RBD)

Tables S4 and S5 provide detailed lists of pathways and path lengths to residue 501 of the U protomer, from residue 614 and from the furin cleavage site (FCS) 685, respectively, of all three protomers. In our calculations, we consider all residues from RBM and FCS. For simplicity, we list the details for these two source/target residues only. The network representation for the case of the G-form in the one-up conformation, as well as the average path lengths, are depicted in Fig. S4

**Figure S4:**
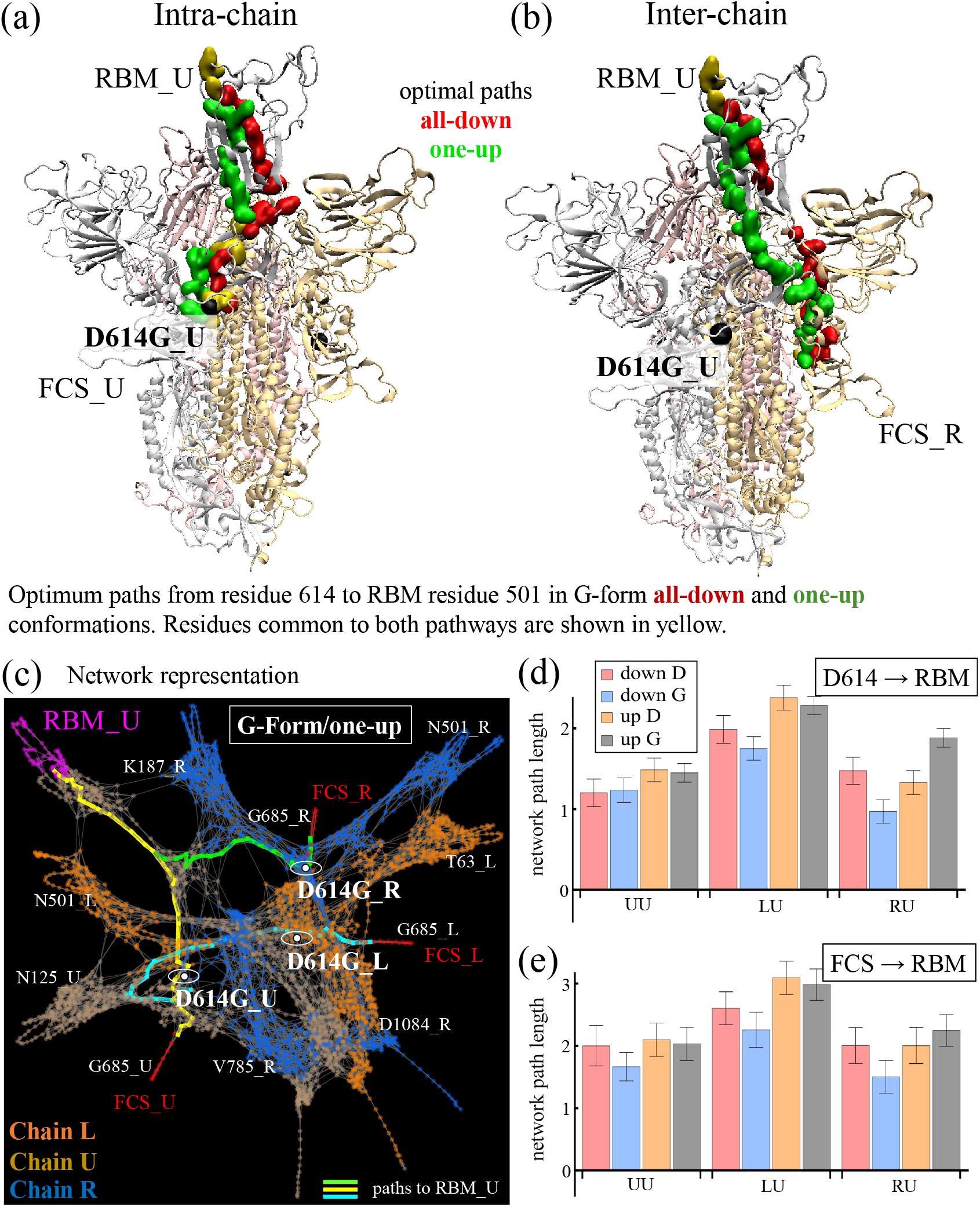
Optimal paths from residue 614 of the U protomer (a) and the R protomer (b) to the RBM residue 501 of the U protomer. (c) Pathways to RMB U in the network representation of the Spike protein from the FCS 685 for each of the protomers (colored paths) for the one-up conformation in the G-Form in the network representation of the Spike protein. Paths pass next residue 614 (white nodes in circles). (d-e) Average path lengths to all residues of the up-RBM from residue 614 (d) and FCS (e) of the three protomers.

**Table S4:** Pathways to residue 501 of the up-RBM from residue 614 of all protomers for the D- and G-form, in the all-down and the one-up conformations.

| source → target | length | path (D-form/all-down) |
| --- | --- | --- |
| D614_L→N501_U | 1.87582 | D614_L, Q613_L, F318_L, R319_L, V320_L, C538_L, V551_L, L552_L, D586_L, A575_L, G566_L, R44_U, Y279_U, K278_U, A288_U, V289_U, S297_U, L296_U, V597_U, S596_U, V595_U, F318_U, R319_U, V320_U, P322_U, N540_U, F541_U, N542_U, N544_U, L390_U, C391_U, F515_U, S514_U, L513_U, V512_U, V511_U, V510_U, R509_U, Y508_U, P507_U, Q506_U, Y505_U, N501_U |
| D614_U→N501_U | 1.09255 | D614_U, F592_U, S591_U, C590_U, G550_U, F541_U, N542_U, N544_U, L390_U, C391_U, F515_U, S514_U, L513_U, V512_U, V511_U, V510_U, R509_U, Y508_U, P507_U, Q506_U, Y505_U, N501_U |
| D614_R→N501_U | 1.37853 | D614_R, Q613_R, Y612_R, L611_R, V610_R, P295_R, D294_R, L293_R, D290_R, V289_R, A288_R, V36_R, Y37_R, Y204_R, I203_R, K202_R, F201_R, Y200_R, E516_U, F515_U, S514_U, L513_U, V512_U, V511_U, V510_U, R509_U, Y508_U, P507_U, Q506_U, Y505_U, N501_U |
| source → target | length | path (D-form/one-up) |
| D614_L→N501_U | 2.23281 | D614_L, Q613_L, G594_L, G593_L, F592_L, C590_L, G550_L, N540_L, F541_L, V327_L, R328_L, P330_L, I332_L, N360_L, S359_L, I358_L, Y396_L, A397_L, D398_L, S399_L, F400_L, V401_L, Y451_L, L452_L, Q493_L, L492_L, F490_L, Y489_L, F377_U, A435_U, Y508_U, P507_U, Q506_U, N501_U |
| D614_U→N501_U | 1.35858 | D614_U, Q613_U, G594_U, G593_U, F592_U, S591_U, C590_U, G550_U, N540_U, F541_U, V327_U, R328_U, S530_U, K529_U, K528_U, P527_U, A363_U, F338_U, F342_U, V511_U, V510_U, R509_U, Y508_U, P507_U, Q506_U, N501_U |
| D614_R→N501_U | 1.20224 | D614_R, Q613_R, V595_R, S596_R, V597_R, T299_R, C301_R, L276_R, L277_R, K278_R, Y279_R, R44_R, G566_U, V576_U, R577_U, D578_U, R328_U, S530_U, K529_U, K528_U, P527_U, A363_U, F338_U, F342_U, V511_U, V510_U, R509_U, Y508_U, P507_U, Q506_U, N501_U |
| source → target | length | path (G-form/all-down) |
| D614G_L→N501_U | 1.71229 | D614G_L, Q613_L, Y612_L, L611_L, V610_L, V597_L, L296_L, S297_L, V289_L, A288_L, I285_L, T284_L, G283_L, R44_L, G566_R, A575_R, D586_R, L552_R, V551_R, C538_R, V320_R, F318_R, N317_R, S316_R, T315_R, V597_R, P295_R, D294_R, C291_R, T274_R, R273_R, Q271_R, L270_R, G89_R, K195_R, F201_R, P230_R, Y396_U, A397_U, D398_U, S399_U, F400_U, V401_U, P507_U, Q506_U, T500_U, N501_U |
| D614G_U→N501_U | 1.15282 | D614G_U, V615_U, F592_U, C590_U, C538_U, V539_U, N540_U, F541_U, V327_U, R328_U, F329_U, P330_U, I332_U, C361_U, S359_U, V395_U, Y396_U, A397_U, D398_U, S399_U, F400_U, V401_U, P507_U, Q506_U, T500_U, N501_U |
| D614G_R→N501_U | 0.930889 | D614G_R, Q613_R, Y612_R, L611_R, V610_R, V597_R, P295_R, D294_R, C291_R, T274_R, R273_R, Q271_R, L270_R, G89_R, K195_R, F201_R, P230_R, Y396_U, A397_U, D398_U, S399_U, F400_U, V401_U, P507_U, Q506_U, T500_U, N501_U |
| source → target | length | path (G-form/one-up) |
| D614G_L→N501_U | 2.17649 | D614G_L, G593_L, F592_L, S591_L, C590_L, V551_L, L552_L, L585_L, V576_L, F565_L, F43_U, R44_U, Y279_U, K278_U, L277_U, L276_U, C301_U, T299_U, Y313_U, Q314_U, T315_U, S316_U, N317_U, F318_U, V320_U, Q321_U, P322_U, N540_U, F541_U, V327_U, S530_U, K529_U, K528_U, N388_U, L387_U, C432_U, V433_U, V511_U, V510_U, R509_U, Y508_U, Q506_U, N501_U |
| D614G_U→N501_U | 1.34378 | D614G_U, Q613_U, Y612_U, V595_U, T315_U, S316_U, N317_U, F318_U, V320_U, Q321_U, P322_U, N540_U, F541_U, V327_U, S530_U, K529_U, K528_U, N388_U, L387_U, C432_U, V433_U, V511_U, V510_U, R509_U, Y508_U, Q506_U, N501_U |
| D614G_R→N501_U | 1.7775 | D614G_R, Q613_R, V595_R, S596_R, Y313_R, T299_R, C301_R, L276_R, L277_R, I285_R, T284_R, L560_U, F559_U, R577_U, D578_U, R328_U, S530_U, K529_U, K528_U, N388_U, L387_U, C432_U, V433_U, V511_U, V510_U, R509_U, Y508_U, Q506_U, N501_U |

**Table S5:** Pathways to residue 501 of the up-RBM from residue 685 of all protomers for the D- and G-form, in the all-down and the one-up conformations.

| source → target | length | path (**D-form/all-down**) |
| --- | --- | --- |
| G685_L→N501_U | 2.63993 | G685_L, S686_L, V687_L, A688_L, S689_L, Q690_L, S691_L, I692_L, A609_L, V610_L, L611_L, Y612_L, F318_L, R319_L, V320_L, C538_L, V551_L, L552_L, D586_L, A575_L, G566_L, R44_U, Y279_U, K278_U, A288_U, V289_U, S297_U, L296_U, V597_U, S596_U, V595_U, F318_U, R319_U, V320_U, P322_U, N540_U, F541_U, N542_U, N544_U, L390_U, C391_U, F515_U, S514_U, L513_U, V512_U, V511_U, V510_U, R509_U, Y508_U, P507_U, Q506_U, Y505_U, N501_U |
| G685_U→N501_U | 2.05054 | G685_U, S686_U, V687_U, A688_U, S689_U, Q690_U, S691_U, I692_U, A609_U, V610_U, L611_U, Y612_U, F318_U, R319_U, V320_U, P322_U, N540_U, F541_U, N542_U, N544_U, L390_U, C391_U, F515_U, S514_U, L513_U, V512_U, V511_U, V510_U, R509_U, Y508_U, P507_U, Q506_U, Y505_U, N501_U |
| G685_R→N501_U | 2.1933 | G685_R, A684_R, G683_R, S682_R, P681_R, S680_R, N679_R, T678_R, T676_R, Q675_R, Y674_R, P600_R, K310_R, E309_R, V308_R, K300_R, L276_R, L277_R, I285_R, A222_R, L223_R, Y204_R, I203_R, K202_R, F201_R, Y200_R, E516_U, F515_U, S514_U, L513_U, V512_U, V511_U, V510_U, R509_U, Y508_U, P507_U, Q506_U, Y505_U, N501_U |
| source → target | length | path (**D-form/one-up**) |
| G685_L→N501_U | 3.13669 | G685_L, S686_L, V687_L, A688_L, S689_L, S691_L, I692_L, A609_L, V610_L, V595_L, G594_L, G593_L, F592_L, C590_L, G550_L, N540_L, F541_L, V327_L, R328_L, P330_L, I332_L, N360_L, S359_L, I358_L, Y396_L, A397_L, D398_L, S399_L, F400_L, V401_L, Y451_L, L452_L, Q493_L, L492_L, F490_L, Y489_L, F377_U, A435_U, Y508_U, P507_U, Q506_U, N501_U |
| G685_U→N501_U | 2.11265 | G685_U, S686_U, V687_U, A688_U, S689_U, Q690_U, S691_U, I692_U, A609_U, V610_U, L611_U, V595_U, F318_U, R319_U, V320_U, Q321_U, P322_U, N540_U, F541_U, V327_U, R328_U, S530_U, K529_U, K528_U, P527_U, A363_U, F338_U, F342_U, V511_U, V510_U, R509_U, Y508_U, P507_U, Q506_U, N501_U |
| G685_R→N501_U | 2.07156 | G685_R, S686_R, V687_R, A688_R, S689_R, S691_R, I692_R, V608_R, L296_R, S297_R, C291_R, L276_R, L277_R, K278_R, Y279_R, R44_R, G566_U, V576_U, R577_U, D578_U, R328_U, S530_U, K529_U, K528_U, P527_U, A363_U, F338_U, F342_U, V511_U, V510_U, R509_U, Y508_U, P507_U, Q506_U, N501_U |
| source → target | length | path (**G-form/all-down**) |
| G685_L→N501_U | 2.24178 | G685_L, S686_L, V687_L, A688_L, S689_L, S691_L, I692_L, V608_L, L296_L, S297_L, V289_L, A288_L, I285_L, T284_L, G283_L, R44_L, G566_R, A575_R, D586_R, L552_R, V551_R, C538_R, V320_R, F318_R, N317_R, S316_R, T315_R, V597_R, P295_R, D294_R, C291_R, T274_R, R273_R, Q271_R, L270_R, G89_R, K195_R, F201_R, P230_R, Y396_U, A397_U, D398_U, S399_U, F400_U, V401_U, P507_U, Q506_U, T500_U, N501_U |
| G685_U→N501_U | 1.79819 | G685_U, S686_U, V687_U, A688_U, S689_U, S691_U, I692_U, I598_U, V597_U, S596_U, V595_U, G594_U, G593_U, F592_U, C590_U, C538_U, V539_U, N540_U, F541_U, V327_U, R328_U, F329_U, P330_U, I332_U, C361_U, S359_U, V395_U, Y396_U, A397_U, D398_U, S399_U, F400_U, V401_U, P507_U, Q506_U, T500_U, N501_U |
| G685_R→N501_U | 1.57206 | G685_R, S686_R, V687_R, A688_R, S689_R, S691_R, I692_R, A609_R, V597_R, P295_R, D294_R, C291_R, T274_R, R273_R, Q271_R, L270_R, G89_R, K195_R, F201_R, P230_R, Y396_U, A397_U, D398_U, S399_U, F400_U, V401_U, P507_U, Q506_U, T500_U, N501_U |
| source → target | length | path (**G-form/one-up**) |
| G685_L→N501_U | 2.99415 | G685_L, S686_L, V687_L, A688_L, S689_L, S691_L, I692_L, A609_L, V610_L, L611_L, Y612_L, F318_L, R319_L, V320_L, C538_L, V551_L, L552_L, L585_L, V576_L, F565_L, F43_U, R44_U, Y279_U, K278_U, L277_U, L276_U, C301_U, T299_U, Y313_U, Q314_U, T315_U, S316_U, N317_U, F318_U, V320_U, Q321_U, P322_U, N540_U, F541_U, V327_U, S530_U, K529_U, K528_U, N388_U, L387_U, C432_U, V433_U, V511_U, V510_U, R509_U, Y508_U, Q506_U, N501_U |
| G685_U→N501_U | 2.03041 | G685_U, S686_U, V687_U, A688_U, S689_U, S691_U, I692_U, A609_U, V610_U, S596_U, T315_U, S316_U, N317_U, F318_U, V320_U, Q321_U, P322_U, N540_U, F541_U, V327_U, S530_U, K529_U, K528_U, N388_U, L387_U, C432_U, V433_U, V511_U, V510_U, R509_U, Y508_U, Q506_U, N501_U |
| G685_R→N501_U | 2.42439 | G685_R, S686_R, V687_R, A688_R, S689_R, S691_R, I692_R, P600_R, K310_R, E309_R, V308_R, F306_R, L48_R, V47_R, R44_R, F43_R, F565_U, V576_U, R577_U, D578_U, R328_U, S530_U, K529_U, K528_U, N388_U, L387_U, C432_U, V433_U, V511_U, V510_U, R509_U, Y508_U, Q506_U, N501_U |

### D. Involvement of the hinge residues of the up-RBD in shortest paths from 614

We explore the role of hinges H_1_ (residues 320-336) and H_2_ (residues 516-536) residues at connecting residue 614 to the RBM employing the Floyd-Warshall algorithm to compute the shortest pathways.

#### Method i

We quantify the involvement of each of the hinges by counting the number of times a residue belonging to a particular hinge is identified to belong to a shortest path (H*_i_* role column). We examine the total set of 854 paths accounting for all the source/target combinations of the three protomers for our four Spike configurations: D form all-down, D form one-up, G form all-down, and G form one-up.

We find that in the all-down configuration there is a marked preference for using H_1_ over H_2_, and using routes that do not pass through the hinges. This finding is more prominent in the D form than the G form. The one-up configuration is characterized by the use of both hinges at comparable amounts with a small preference towards H_1_, and a highly more frequent use of the hinge residues in the G form than in the D form. Finally, the use of H_1_ occurs more frequent for intra-protomer pathways, while H_2_ is found more often for inter-protomer pathways.

D form / all-down

| source | target | $H_1$ role | residues | $H_2$ role | residues |
| --- | --- | --- | --- | --- | --- |
| D614_U | RBM_U | 0 | - | 0 | - |
| D614_L | RBM_U | 198 | V320_L, V320_U, P322_U | 0 | - |
| D614_R | RBM_U | 0 | - | 24 | E516_U |

D form / one-up

| source | target | $H_1$ role | residues | $H_2$ role | residues |
| --- | --- | --- | --- | --- | --- |
| D614_U | RBM_U | 142 | V327_U, R328_U | 284 | P527_U, K528_U, K529_U, S530_U |
| D614_L | RBM_U | 284 | V327_L, R328_L, P330_L, I332_L | 0 | - |
| D614_R | RBM_U | 71 | R328_U | 284 | P527_U, K528_U, K529_U, S530_U |

G form / all-down

| source | target | $H_1$ role | residues | $H_2$ role | residues |
| --- | --- | --- | --- | --- | --- |
| D614G_U | RBM_U | 280 | V327_U, R328_U, F329_U, P330_U, I332_U | 0 | - |
| D614G_L | RBM_U | 71 | V320_R | 0 | - |
| D614G_R | RBM_U | 0 | - | 0 | - |

G form / one-up

| source | target | $H_1$ role | residues | $H_2$ role | residues |
| --- | --- | --- | --- | --- | --- |
| D614G_U | RBM_U | 284 | V320_U, Q321_U, P322_U, V327_U | 213 | K528_U, K529_U, S530_U |
| D614G_L | RBM_U | 284 | V320_U, Q321_U, P322_U, V327_U | 213 | K528_U, K529_U, S530_U |
| D614G_R | RBM_U | 71 | R328_U | 213 | K528_U, K529_U, S530_U |

#### Method ii

To complement the analysis, we consider the role of suboptimal residues and whether or not hinge residues can be present withing this group. To this end, we count the hinge residues in the top-50 suboptimal group connecting 614 with RBM. As a general rule, it is found that the usage of H_1_ is more prominent than H_2_, especially in the D form. Also, the all-down configuration tends to use H_1_ residues on a greater proportion than H_2_ residues. The suboptimal residues are identified by comparing the direct pathlength from 614 and RBM with an indirect pathway that go through an additional third node. The difference in the length of these pathways provides a score of how good this third node is at connecting 614 and RBM through a suboptimal path.

D form / all-down

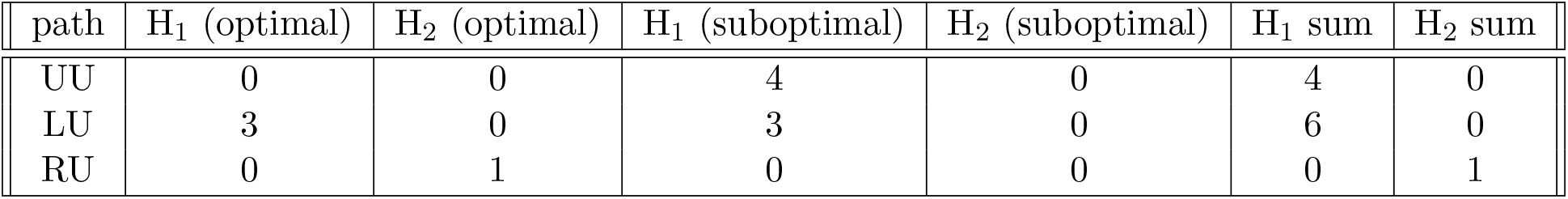

| path | $H_1$ (optimal) | $H_2$ (optimal) | $H_1$ (suboptimal) | $H_2$ (suboptimal) | $H_1$ sum | $H_2$ sum |
| --- | --- | --- | --- | --- | --- | --- |
| UU | 0 | 0 | 4 | 0 | 4 | 0 |
| LU | 3 | 0 | 3 | 0 | 6 | 0 |
| RU | 0 | 1 | 0 | 0 | 0 | 1 |

D form / one-up

| path | $H_1$ (optimal) | $H_2$ (optimal) | $H_1$ (suboptimal) | $H_2$ (suboptimal) | $H_1$ sum | $H_2$ sum |
| --- | --- | --- | --- | --- | --- | --- |
| UU | 2 | 4 | 7 | 7 | 9 | 11 |
| LU | 4 | 0 | 7 | 0 | 11 | 0 |
| RU | 1 | 4 | 3 | 1 | 4 | 5 |

G form / all-down

| path | $H_1$ (optimal) | $H_2$ (optimal) | $H_1$ (suboptimal) | $H_2$ (suboptimal) | $H_1$ sum | $H_2$ sum |
| --- | --- | --- | --- | --- | --- | --- |
| UU | 5 | 0 | 3 | 7 | 8 | 7 |
| LU | 1 | 0 | 2 | 0 | 3 | 0 |
| RU | 0 | 0 | 0 | 0 | 0 | 0 |

G form / one-up

| path | $H_1$ (optimal) | $H_2$ (optimal) | $H_1$ (suboptimal) | $H_2$ (suboptimal) | $H_1$ sum | $H_2$ sum |
| --- | --- | --- | --- | --- | --- | --- |
| UU | 4 | 3 | 6 | 7 | 10 | 10 |
| LU | 4 | 3 | 5 | 3 | 9 | 6 |
| RU | 1 | 3 | 2 | 2 | 3 | 5 |

### E. Impact of residue modification/deletions in the protein network

The relevance of a particular node in a network can change with the graph’s topology. In the main manuscript, we showed that the closeness centrality of the NTD residues modified/deleted by the Delta variant, on average, tend to score higher than that of the NTD supersite. Here we report the centrality scores of all modifications/deletions characteristic of the Delta variant, as well as, the deletions reported in the Alpha and Beta variants [2]. In the latter we focus in deletions since in the Alpha variant there are no substitutions in the NTD, only deletions, and for the Beta variant, we find that the deleted residues score higher than the substitutions in the NTD.

Figure S5 illustrate our finding of betweenness and eigenvector centrality ranking of the modified/deleted residues in the Delta variant compared to the rest of the residues in the protein. It is found that modified/deleted residues in Delta rank very low in these two metrics. A similar calculation but for closeness centrality shows a different story where these critical residues occupy more dominant positions, especially those belonging to the NTD. Figure S6(a) illustrates this finding for the all-down configuration of the Spike protein. A comparison of the closeness centrality of the RBD sites modified by the Delta variant, scores very similar to the RBD site associated to high-frequency mutations (sites 417, 439, 452, 484, 501) [4, 5, 6]. [Fig. S5(b)].

**Figure S5:**
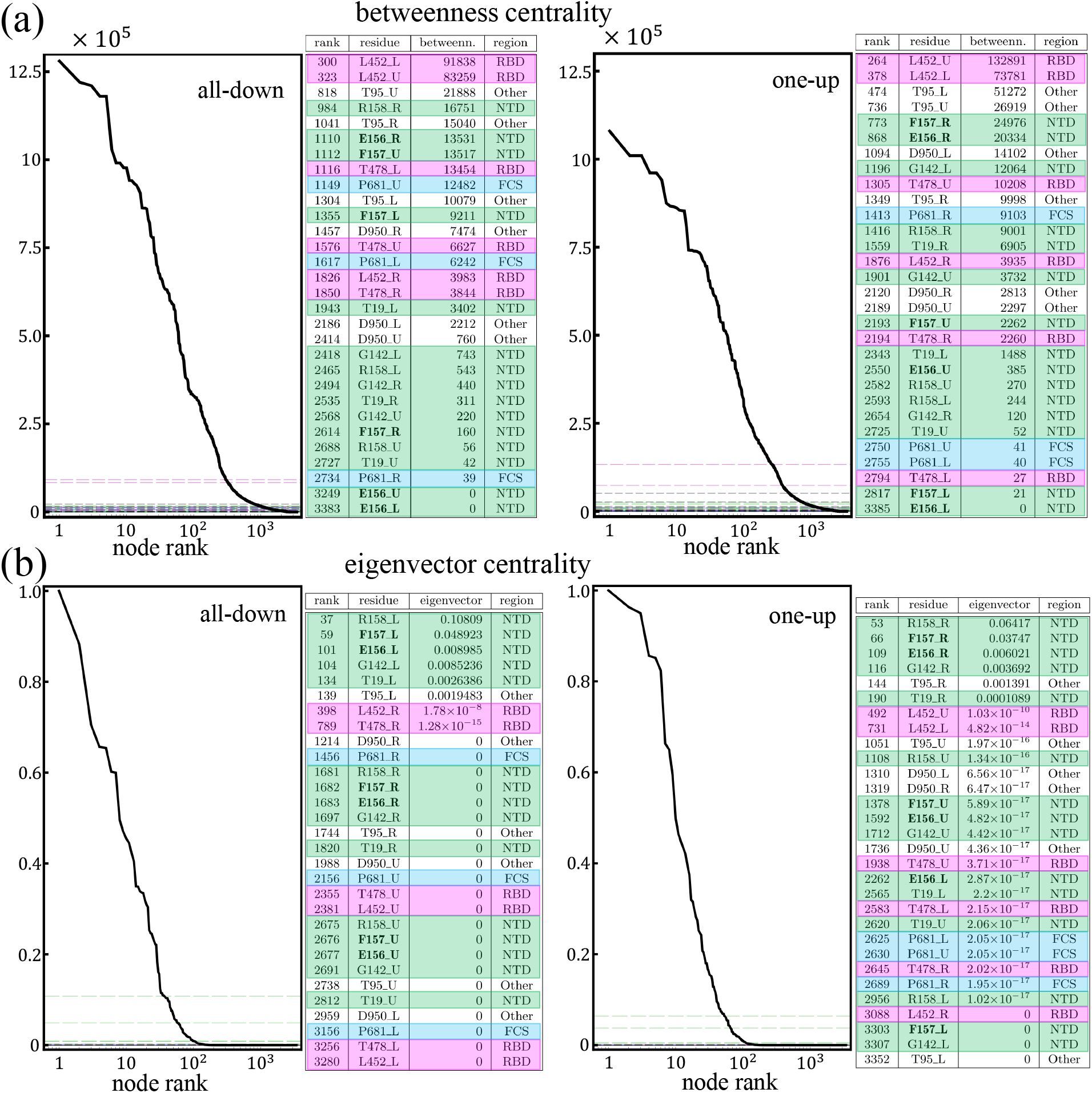
Betweenness (a) and eigenvector centrality (b) of residues modified by the Delta variant. Each panel shows the ranking of these residues (horizontal grid lines) within the whole protein, and a table that details specific residue information (rank, residue identity, centrality score, and protein region where the residue belongs to), for the all-down (left panel) and the one-up (right panel). Colors in the gid lines of the centrality plots and the tables are assigned by the protein region each residue belongs to: pink for RBD, green for NTD supersite, blue for FCS, and white for other regions.

**Figure S6:**
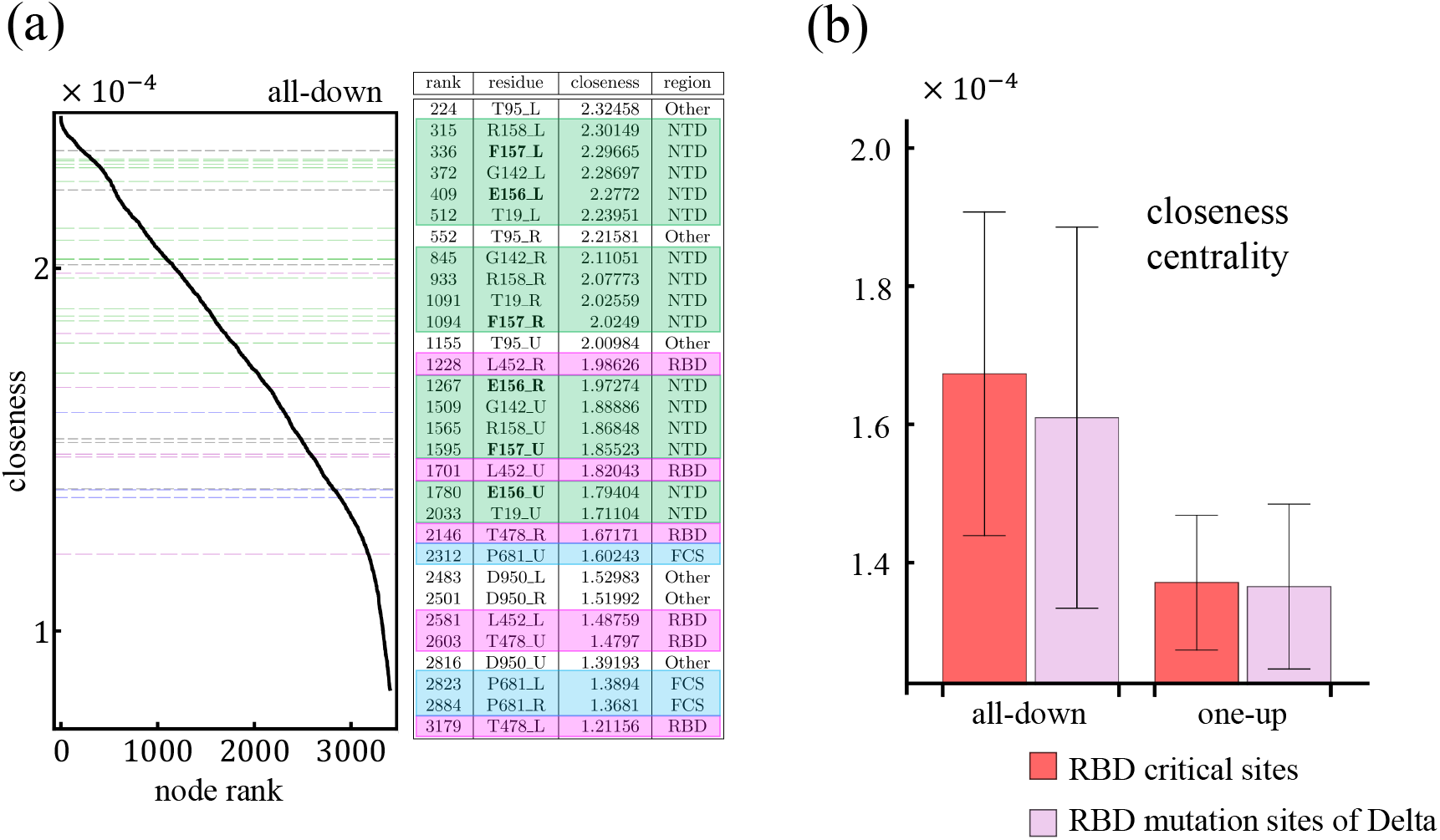
(a) Closeness centrality of residues modified by the Delta variant for the all-down conformation of the Spike protein. The left panel shows the ranking of these residues (horizontal grid lines) within the whole protein, and a table that details specific residue information (rank, residue identity, centrality score, and protein region where the residue belongs to), for the all-down (left panel) and the one-up (right panel). Colors in the gid lines of the centrality plots and the tables are assigned by the protein region each residue belongs to: pink for RBD, green for NTD supersite, blue for FCS, and white for other regions. (b) Average closeness centrality of the residues in the RBD modified by the Delta variant compared to the RBD sites with high frequency mutations (sites 417, 439, 452, 484, 501).

Examining the impact of the NTD residues altered in the Alpha, Beta, and Gamma variants of SARS-CoV-2 Spike protein. Some of these residues include deletions, reported to occur in the NTD domain, that are linked to antibody escape and they tend to overlap with the NTD supersite [3] [Fig. S7(a)]. Using the data from Los Alamos National Laboratory Sequence Entropy Data obtained from cov.lanl.gov database, as of July 22, 2021, the Alpha variant is characterized by the deletion of residues 69-70, and 144, the Beta variant the mutated residues in the NTD are 18, 80, 215, 242-244, and the Gamma variant the mutations in the NTD are 18,20,26,138,190. Comparing the average closeness centrality of these deleted residues with that of the NTD supersite, we find that the score of the mutations associated to the Gamma and Beta variants rank higher [Fig. S7(b)]. The scores of the residues deleted by the Alpha variant and those of the NTD supersite are very similar to each other, though the former is slightly larger than the latter. The ranking of these residues within the whole protein is shown in Fig. S7(c) revealing that these key residues occupy dominant position in the protein ranking in closeness centrality. These residues are very efficient at initiating communication pathways capable to reach the whole protein.

A computation of the betweenness and eigenvector centralities of the residues deleted by the Alpha and Beta variants, shows that these residues rank very low in these metrics within the entire protein [Fig. S7]. For betweenness, we find that the impact of these residues often changes depending on which conformation we are looking at: the all-down or the one-up. For example, residu<u>e</u> L242_C (deleted in the Beta variant) in the down conformation ranks in the top 8% of most critical nodes in terms of communication (i.e., betweenness centrality). However, in the up conformation this residue is no longer relevant carrying a value of betweenness equal to zero. Another example is residue H69_A (in Alpha), which in the down conformation, it helps linking three pairs of nodes, while in the up configuration, it is found to take part in the shortest path of nearly seven thousand pairs of nodes. Therefore, deletion of these two residues is likely to affect severely only one of the conformations. On the other hand, the deletion of residues L242 A, A243, and L244 (all three in Beta), is expected to impact the protein the most because they hold large values of betweenness centralities for both conformations. For the all-down G, the most relevant nodes associated to deletion mutations are L242 C, A243 C, L244 C, L242 A (Beta). These nodes hold betweenness centrality values within the top 8% among all of the residues of the Spike. For the one-up G, residue A243 B, A243 A, A243 C, L242 A hold values of betweenness in the top 12% among all of the residues in the Spike.

**Figure S7:**
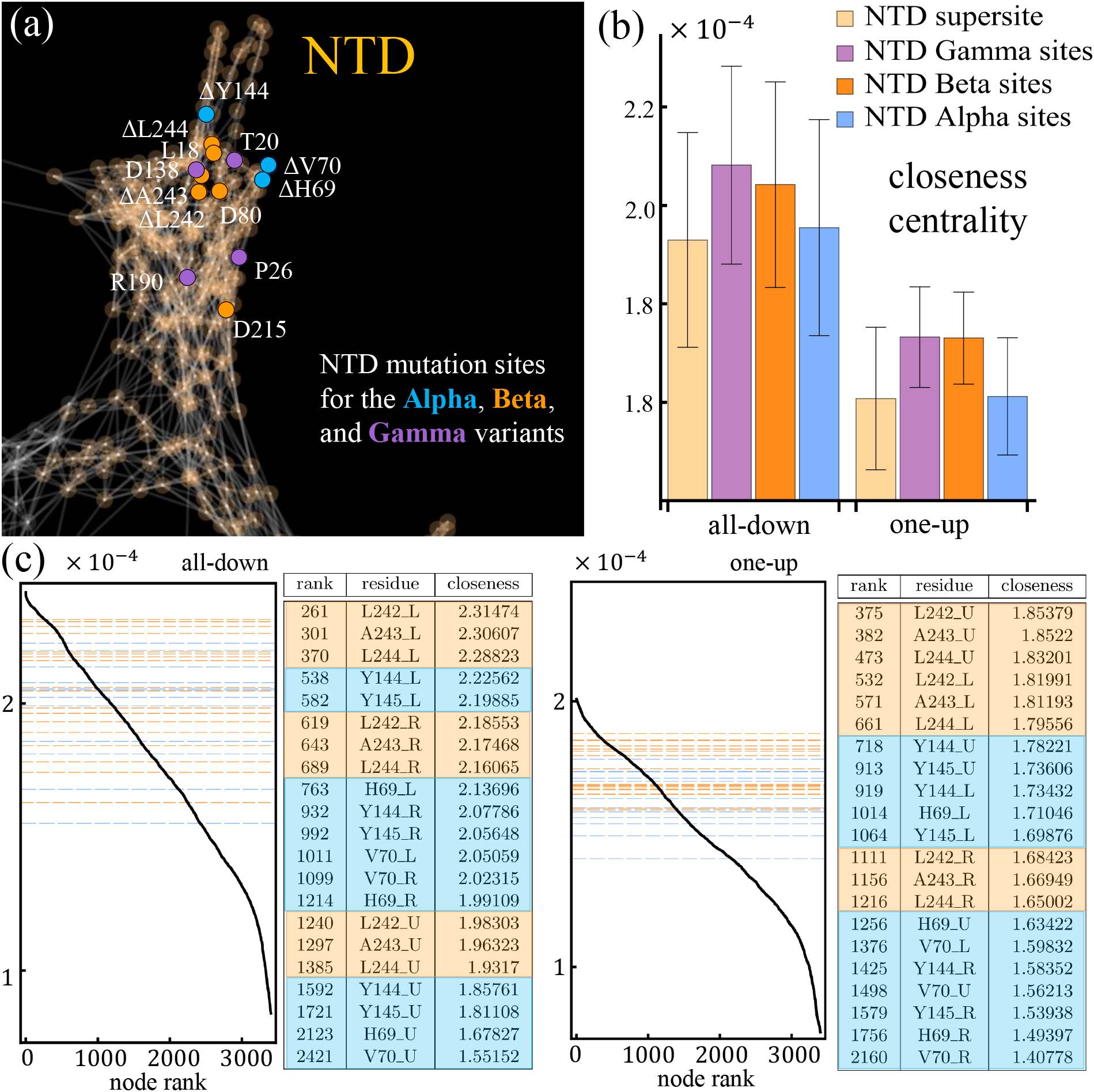
(a) Spike protein in the network representation highlighting the location of modified/deleted residues in the Alpha (blue), Beta (orange), and Gamma (purple) variants of the SARS-CoV-2 virus. (b) Comparison of the average closeness centrality of residues deleted by the Alpha (blue), Beta (orange), and Gamma (purple), and that of the NTD supersite (light orange). (c) Ranking of the closeness centrality for these deleted residues (horizontal grid lines) within the whole protein, and a table that details specific residue information (rank, residue identity, and centrality score times 10^4^), for the all-down (left panel) and the one-up (right panel). Colors in the gid lines of the centrality plots and the tables are assigned by the variant: blue for Alpha, and orange for Beta.

**Figure S8:**
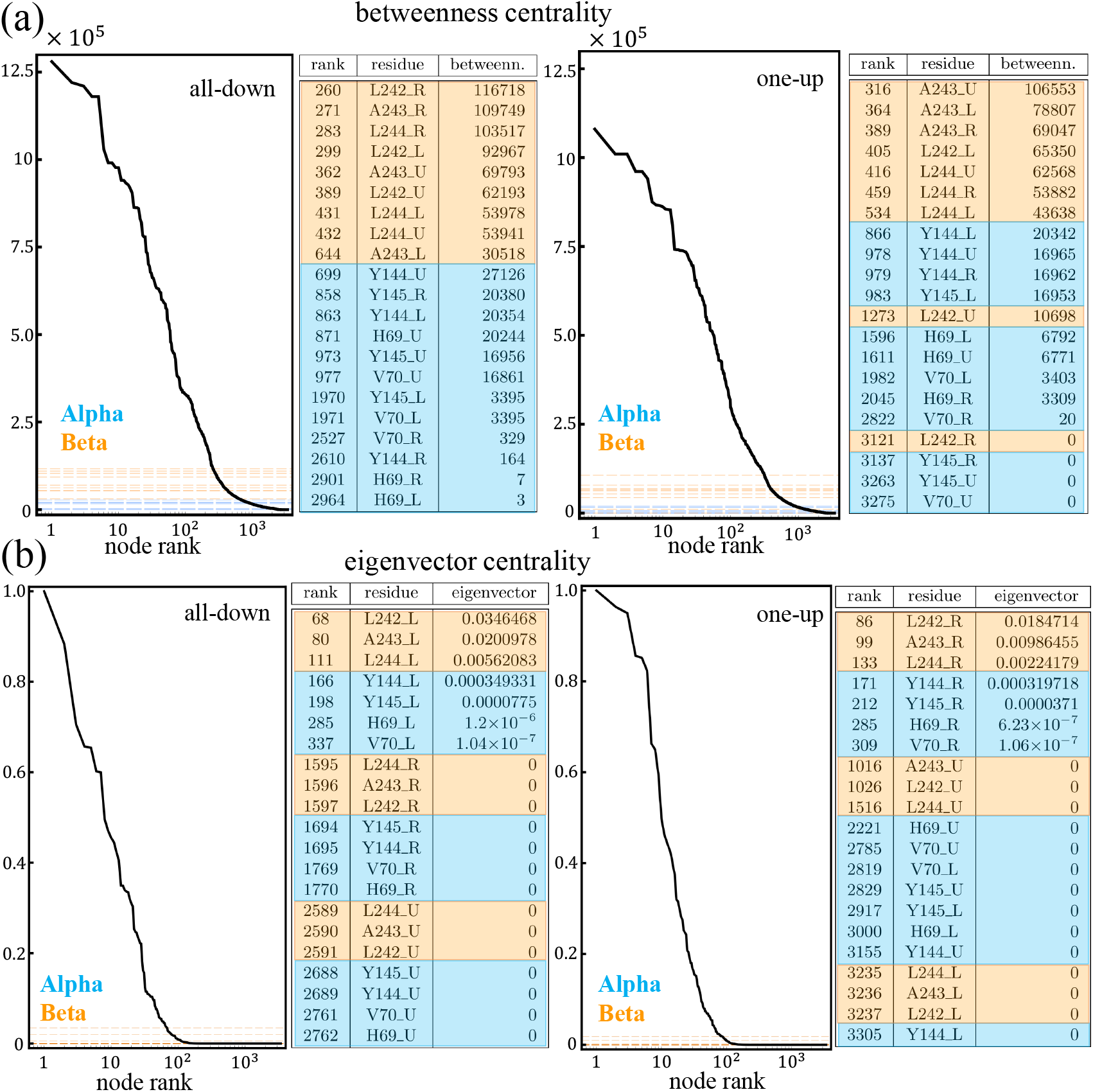
Betweenness (a) and eigenvector centrality (b) of deleted residues of the Alpha (blue) and Beta (variant). Each panel shows the ranking of these residues (horizontal grid lines) within the whole protein, and a table that details specific residue information (rank, residue identity, and centrality score), for the all-down (left panel) and the one-up (right panel).

### F. High frequency mutable residues

Conservation entropy measures the mutation frequency associated to each residue. Table S6 shows the entropy for the highest mutable residues of the Spike protein alongside with their respective betweenness centrality from our network analysis. Residues ranking below this top-60 in conservation entropy, hold entropy values lower than the 1% of the highest ranked residue in conservation entropy. These high-frequency mutable residues do not overlap with the top 10% of high betweenness residue ranks. Data from Los Alamos National Laboratory Sequence Entropy Data obtained from cov.lanl.gov database, as of July 22, 2021.

### G. Harnessing centrality measures to characterize functionality regions in the N-terminal domain

Finally, we look into how based on the centrality measures and average residue exposure, we are able to distinguish two constrasting regions within the NTD: one region (region 1) where a large number of mutations and epitopes are found, and a second one (region 2) that is potentially relevant for functionality and where, up to August 2021, no mutations have been found. We identify region 1 by looking into residues with low-betweenness (< 2.15% of the highest), low-eigenvector (< 9% of the highest), mid-closeness (between 71 and 92% of the highest), and high-exposure (> 16% of exposure). Total exposure of each residue is obtained from theoretical surface area calculations of Gly-X-Gly tripeptides [7]. This criteria identfies 80.4% of the residues belonging to the NTD super site, as well as, 63.1% of the NTD residues altered by the Alpha, Beta, Gamma, and Delta variants. On the other hand, region 2 is characterized by sole condition of having eigenvector centrality scores higher than 12% of the highest. This single condition carries some interesting implications for the other centrality measures. For example, we find that residues in region 2 hold a very uniform closeness centrality values that lie between 91 and 94% of the highest. Similarly, we find that these residues hold vertex strength values higher than 25% of the highest. These findings become the first step towards predicting where the next mutations could take place and hence talk about pandemic preparedness. Figure S9 Illustrates our finding for these constrasting regions.

**Table S6:** Top 60 highest mutable residues according to their conservation entropy score. Residues below this ranking have entropy scores below the 1% of the highest ranked residue in conservation entropy. Data from Los Alamos National Laboratory Sequence Entropy Data server cov.lanl.gov database, as of July 22, 2021.

| residue number | residue | conservation entropy | maximum betweenness | rank |
| --- | --- | --- | --- | --- |
| 17 | N | 0.23209 | 10276 | 633 |
| 18 | L | 0.24245 | 11147 | 621 |
| 19 | T | 0.22808 | 6905 | 720 |
| 20 | T | 0.13622 | 34 | 1045 |
| 26 | P | 0.13134 | 16691 | 528 |
| 67 | A | 0.03743 | 13550 | 573 |
| 69 | H | 0.69958 | 6792 | 735 |
| 70 | V | 0.70861 | 3403 | 862 |
| 75 | G | 0.02566 | 6770 | 746 |
| 80 | D | 0.08328 | 6820 | 726 |
| 95 | T | 0.22285 | 51272 | 282 |
| 98 | S | 0.08466 | 6752 | 748 |
| 138 | D | 0.16105 | 4798 | 809 |
| 142 | G | 0.21869 | 12064 | 608 |
| 143 | V | 0.02433 | 36725 | 339 |
| 144 | Y | 0.70251 | 20342 | 467 |
| 152 | W | 0.15643 | 6792 | 736 |
| 153 | M | 0.03797 | 10183 | 641 |
| 156 | E | 0.21532 | 20334 | 468 |
| 157 | F | 0.25236 | 24976 | 420 |
| 158 | R | 0.21555 | 9001 | 677 |
| 188 | N | 0.11426 | 5412 | 793 |
| 190 | R | 0.11518 | 41907 | 321 |
| 215 | D | 0.05308 | 6718 | 755 |
| 222 | A | 0.30369 | 121492 | 184 |
| 242 | L | 0.03616 | 65350 | 242 |
| 243 | A | 0.04127 | 106553 | 199 |
| 244 | L | 0.03592 | 62568 | 248 |
| 253 | D | 0.09903 | 10159 | 648 |
| 262 | A | 0.04366 | 26901 | 403 |
| 272 | P | 0.03329 | 1027 | 990 |
| 417 | K | 0.14337 | 22 | 1054 |
| 439 | N | 0.06871 | 3392 | 878 |
| 452 | L | 0.29757 | 132891 | 168 |
| 477 | S | 0.14296 | 6835 | 725 |
| 478 | T | 0.25073 | 10208 | 637 |
| 484 | E | 0.23143 | 3396 | 873 |
| 494 | S | 0.02955 | 4545 | 821 |
| 501 | N | 0.70502 | 3322 | 889 |
| 570 | A | 0.69536 | 2005 | 945 |
| 583 | E | 0.02634 | 4236 | 829 |
| 614 | D | 0.07355 | 4322 | 825 |
| 655 | H | 0.12143 | 45 | 1039 |
| 675 | Q | 0.06103 | 29387 | 383 |
| 677 | Q | 0.09388 | 12583 | 595 |
| 681 | P | 0.88489 | 9103 | 676 |
| 688 | A | 0.02974 | 22867 | 442 |
| 701 | A | 0.09993 | 12471 | 598 |
| 716 | T | 0.69338 | 161291 | 134 |
| 732 | T | 0.07145 | 7680 | 707 |
| 769 | G | 0.0395 | 6974 | 719 |
| 845 | A | 0.0245 | 6797 | 733 |
| 859 | T | 0.04505 | 12271 | 602 |
| 936 | D | 0.02929 | 3389 | 879 |
| 939 | S | 0.02471 | 4604 | 817 |
| 950 | D | 0.24324 | 14102 | 562 |
| 957 | Q | 0.03744 | 31763 | 367 |
| 982 | S | 0.69182 | 1381 | 979 |
| 1027 | T | 0.12039 | 50658 | 286 |
| 1118 | D | 0.698 | 68693 | 233 |

**Figure S9:**
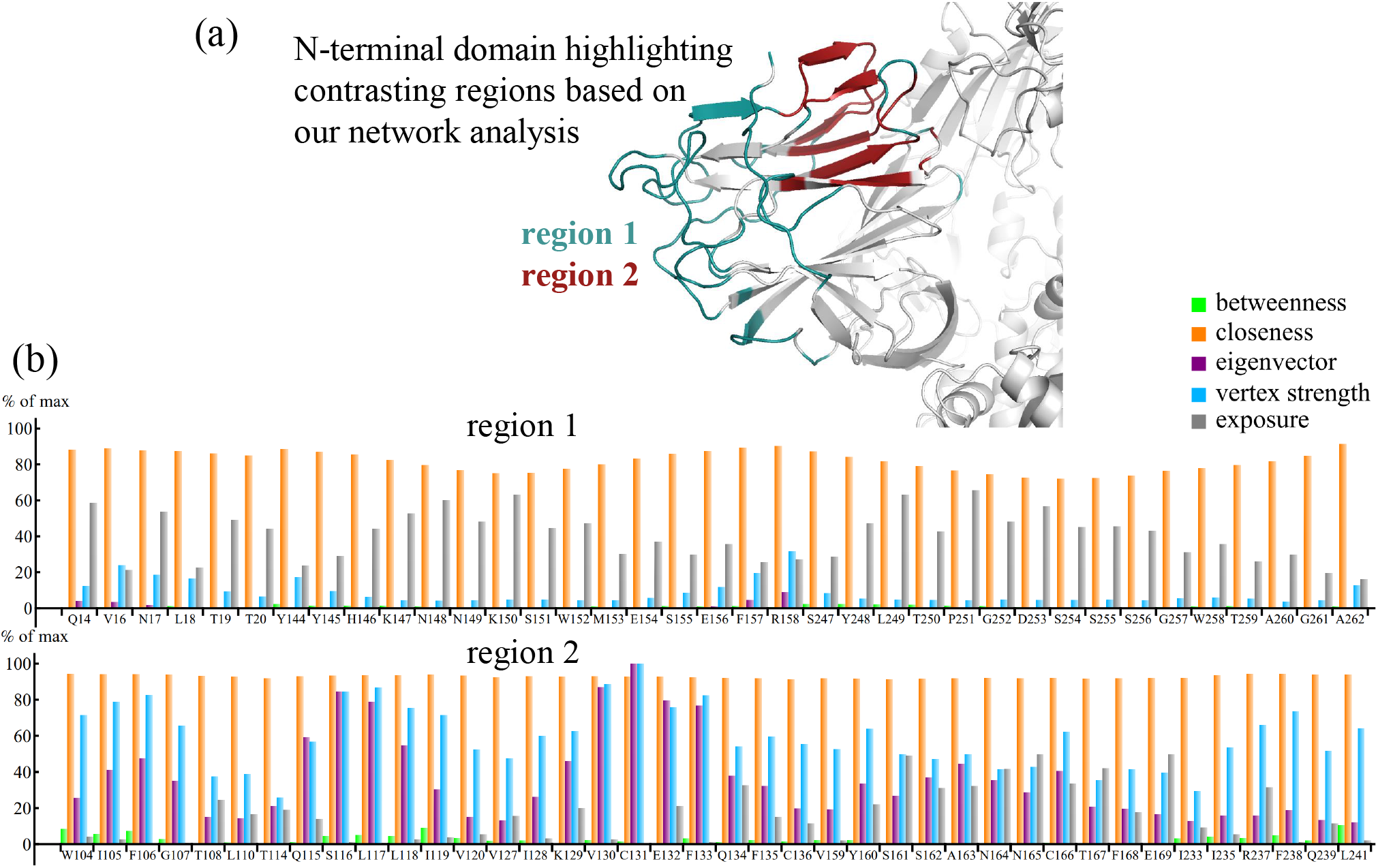
(a) N-terminal domain of the SARS-CoV-2 Spike protein highlighting the regions identified by our network analysis. Region 1 is characterized by specific conditions of low-betweenness, mid-closeness, low-eigenvector, and high-exposure (see text for details), while region 2 follows the sole condition of high eigenvector centrality. (b) Average centrality values of each residue in region 1 (top panel), and region 2 (bottom panel) measured as a porcentage of the highest average centrality found.

